# Dynamic epigenomic landscape of carbon-concentrating mechanisms in the model industrial oleaginous microalga *Nannochloropsis oceanica*

**DOI:** 10.1101/2024.09.06.611585

**Authors:** Yanhai Gong, Qintao Wang, Li Wei, Lianhong Wang, Nana Lv, Xuefeng Du, Chen Shen, Yi Xin, Luyang Sun, Jian Xu

## Abstract

Despite their ecological and physiological significance, how carbon-concentrating mechanisms (CCM) are regulated in microalgae remains elusive. Here in the model industrial microalga *Nannochloropsis oceanica*, we uncovered an epigenetic regulatory mechanism for CCM via comprehensive, multi-dimensional epigenomic analyses. Our integrated study reveals the complex interplay among histone modifications, dynamic nucleosome positioning, and 3D chromatin structure in regulating gene expression during low CO_2_ adaptation, despite minimal DNA methylation. Histone modifications, including lysine acetylation (H3K9ac and H3K27ac), crotonylation (Kcr), and methylation (H3K4me2), were associated with active chromatin states. Significantly altered ChIP-Seq peaks were linked to 43.1% of the differentially expressed genes (DEGs). Notably, H3K4me2 exhibited a distinct dual-peak profile around the transcription start site (TSS), which is unique among microalgae and plants. Chromatin compartment dynamics were correlated with gene expression and histone modifications, particularly H3K4me2, while differentially positioned nucleosomes were associated with key CCM-related genes and transcription factors. To further elucidate the role of H3K4me2, we knocked out its methyltransferase, resulting in genome-wide H3K4me2 peak shifts, slower growth, and reduced photosynthesis. These changes were accompanied by differential expression of key genes of NoHINT and NoPMA2, whose subsequent deletion and overexpression revealed their subtle yet significant impacts on growth and photosynthetic efficiency under low CO_2_ conditions, with NoHINT regulating growth and NoPMA2 influencing photosynthesis. Finally, we proposed a comprehensive model for epigenetic regulation of CCM in *N. oceanica*, which established a foundation for enhancing microalgal productivity through targeted epigenetic modifications.

**Highlights:**

- Multi-layered epigenetic modifications contribute to the regulation of CCM and its adaptation to low CO_2_ in *N. oceanica*.
- The histone modification H3K4me2 regulates the growth and photosynthesis of *N. oceanica* under low CO_2_ conditions.
- H3K4me2 targets NoHINT and NoPMA2 in *N. oceanica*, with NoHINT influencing growth dynamics and NoPMA2 modulating photosynthetic efficiency under low CO_2_ conditions.

## Introduction

Marine microalgae play a crucial role in mitigating atmospheric CO_2_ levels through carbon assimilation, thereby significantly contributing to the reduction of global warming and ocean acidification (Doney, 2010; Hansen et al., 2006). This carbon assimilation process is catalyzed by the bifunctional enzyme ribulose-1,5-bisphosphate carboxylase/oxygenase (Rubisco), which initiates the carbon-fixing Calvin cycle under high CO_2_ conditions but also leads to wasteful photorespiration when the O_2_ to CO_2_ concentration ratio is relatively high (Hayer-Hartl and Hartl, 2020). Since Rubisco’s oxygenase activity (photorespiration) competes with carboxylation, it is crucial for oxygenic photosynthetic organisms to enhance CO_2_ concentration around Rubisco to optimize carbon fixation. This process is particularly important for aquatic plants and microalgae due to the relatively low concentration of dissolved CO_2_ in water, resulting from its limited solubility and slower diffusion rate. Consequently, these organisms rely more heavily on CO_2_-concentrating mechanisms (CCMs) to effectively convert and concentrate CO_2_ around Rubisco for efficient carbon fixation (Bauwe et al., 2010; Hagemann and Bauwe, 2016).

These microalgal CCMs, which are essential for overcoming the limitations of CO_2_ diffusion and the low substrate specificity of Rubisco (Poudel et al., 2020), actively pump exogenous inorganic carbon (Cᵢ) into cells, resulting in the accumulation of CO_2_ at carboxylation sites. Past works from transcriptional (Fang et al., 2012; Wei et al., 2019a), biophysical (Fei et al., 2022; Long et al., 2021), and biochemical (Lin et al., 2020; Wang et al., 2014) perspectives have revealed the complexity and lineage-specificity of the CCM pathway among various microalgal species. Efforts in cyanobacteria and green algae have identified key genes and proteins involved in CCM, including carbonic anhydrases (e.g., CAH1-9), bicarbonate transporters (e.g., HLA3, LCI1, LCIA, CCP1, CCP2, CIA8, BST1-3), and regulatory proteins (e.g., CIA5/CCM1, LCR1) (Kupriyanova et al., 2023; Santhanagopalan et al., 2021; Yoshioka et al., 2004). However, the comprehensive regulatory network governing these processes under varying CO_2_ levels has yet to be fully elucidated, partially owing to the limited understanding of the epigenome in the microalgal CCM, which has been reported to be essential in the regulation of C_4_ and CAM photosynthesis in terrestrial higher plants (Heimann et al., 2013, Shi et al., 2021).

Epigenomic factors such as histone modifications, nuclear positioning, and 3D conformation structures of chromatin work together to govern genomic activity without altering the DNA sequence. In various oxygenic photosynthetic organisms, chromatin states dynamically respond to environmental changes and play essential roles in gene regulation and environmental adaptation (Bacova et al., 2020; Bhadouriya et al., 2021; Ferrari et al., 2023; Gao et al., 2016; Probst and Mittelsten Scheid, 2015). In *Chlamydomonas reinhardtii*, promoter histone modifications and DNA 6mA methylation can reflect the transcriptional status of genes, identifying regulatory switch genes under environmental stress (Fu et al., 2015; Ngan et al., 2015). However, pinpointing causative effects between epigenetic variance, gene expression, and the corresponding phenotype is challenging and may be biased by secondary responses. For example, initial findings in rice suggested that phosphate starvation induces stable hypermethylation near active genes, a counterintuitive result as hypermethylation typically represses gene activity. Further investigation revealed that this hypermethylation acts as a protective response to silence the transposable elements close to the active genes, as a local increase in transcription may lead to reactivation of the transposable element (Secco et al., 2015). Therefore, comprehensive genome-wide epigenetic profiles and experimental validation are needed to identify the causality of epigenomic changes in relation to stress treatments.

*Nannochloropsis* spp. are unicellular oleaginous microalgae known for their high photoautotrophic biomass productivity, elevated lipid content, and suitability for industrial cultivation (Radakovits et al., 2012). *Nannochloropsis* spp. possess a unique CCM characterized by the absence of microcompartments such as carboxysomes or pyrenoids, which are the typical sites for Rubisco and CCM localization in cyanobacteria and green algae (Gee and Niyogi, 2017; Rae et al., 2013; Wang et al., 2015). We showed that 32% of genes in *N. oceanica* were transcriptionally regulated under low CO_2_ stress (Wei et al., 2019a), and dynamics in a few histone marks were also found (Liu and Wei, 2023; Wei and Xu, 2018). However, elucidating the potential roles of epigenetic regulation has been hindered by the lack of a comprehensive epigenomic map.

Here, we generated an integrated epigenome roadmap including histone modifications, nucleosome occupancy and 3D chromatin conformation in *N. oceanica* both before and after the transition to low CO_2_ conditions, and proposed a comprehensive CCM model of epigenetic mechanisms in *N. oceanica* IMET1. We showed that H3K4me2 plays a crucial role in regulating carbon assimilation by targeting key genes such as NoHINT and NoPMA2. The functional validation of these targets through knockout and overexpression demonstrated their subtle yet significant impacts on growth and photosynthetic efficiency. These results highlight the intricate epigenetic control of carbon metabolism in *N. oceanica*, providing new insights into its CCM and potential for enhancing microalgal productivity through targeted epigenetic modifications.

## Results

### Comprehensive Profiling of the Epigenomic Landscape Under Low CO_2_ Stress

We recently showed that *N. oceanica* adapts to low CO_2_ stress by activating CCM while modulating global metabolic networks, including upregulation of photorespiration and tetrahydrofolate cycle, transient upregulation of Calvin cycle and light harvesting genes, downregulation of DNA/RNA metabolism, gene expression, protein synthesis and modification, and transient downregulation of TCA cycle (Wei et al., 2019a). However, the chromatin changes that drive differential gene expression (and thus contribute to low CO_2_ adaption) are unknown.

To understand how genes are regulated during low CO_2_ adaptation, we constructed a multi-dimensional epigenomic atlas from cultures of *N. oceanica* IMET1 that transitioned from high CO_2_ (HC; 5% CO_2_) to low CO_2_ (LC; 0.01% CO_2_) (**Fig. 1**). Microalgal cells were harvested 24 hours post-transition to LC to mitigate confounding effects from circadian rhythms, given that CCM activation occurs within three hours (**Methods**). This comprehensive atlas encompasses genome-wide DNA methylation (5mC), histone modification marks, nucleosome occupancy, and 3D chromatin structure (**Fig. 1**), thus providing an integrated landscape of epigenomic modifications under HC and LC conditions. Special attention was given to correlating these epigenetic changes with gene expression patterns and elucidating the interrelationships among each epigenetic factor. A substantial compendium of 73 datasets comprising 504 gigabases of sequencing data (**Table 1**) was generated (deposited in the Sequence Read Archive database) and integrated into the NanDeSyn database (Gong et al., 2020), providing a valuable resource for microalgal epigenetic research.

**Figure 1.**
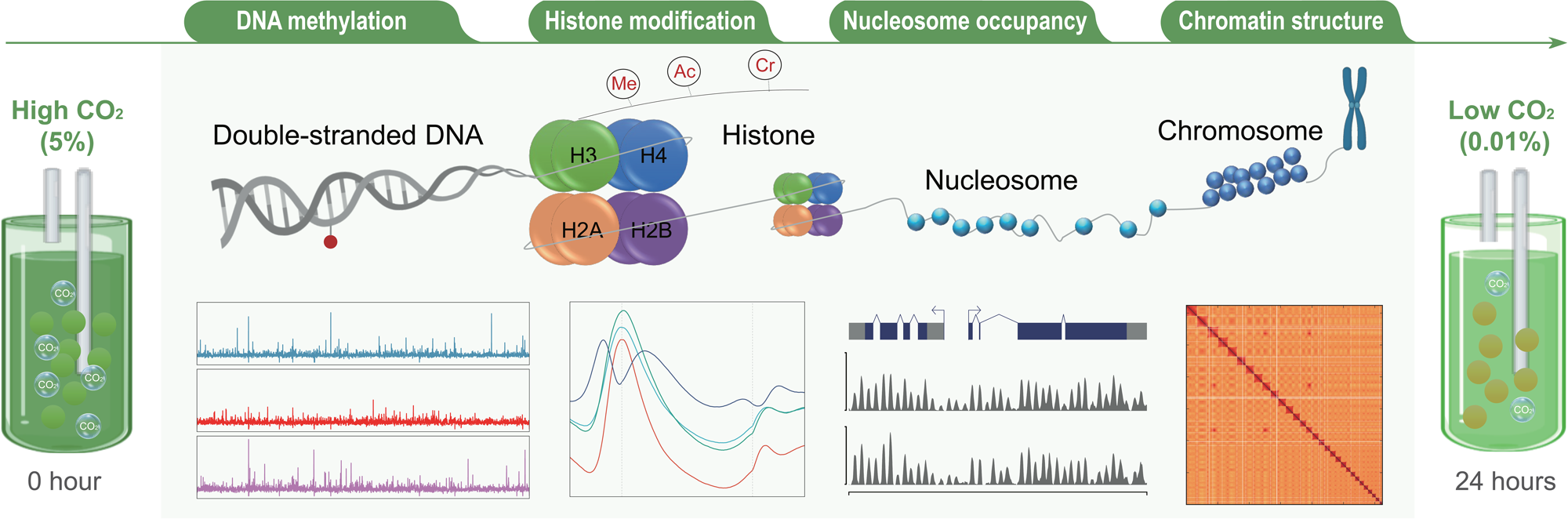
Overview of multi-dimensional epigenomic experiments to dissect the regulatory mechanisms of CCMs in *N. oceanica*. Under low-CO_2_ stress (HC_0h vs LC_24h), histone modifications H3K4me2, H3K9ac, H3K27ac, and Kcr were profiled via ChIP-Seq; nucleosome occupancy was measured via MNase-Seq; chromosome 3D structure was measured via Hi-C; and DNA 5mC methylation was profiled via WGBS. Combined with previous sequenced transcriptome data, how those epigenetic marks regulate gene transcription and determine microalgal phenotypes were investigated.

**Table 1.** The 73 functional-genomics datasets generated as part of this study.

| Experiment | Cell type | Condition | Enzyme/antibody |
| --- | --- | --- | --- |
| **DNA methylation (WGBS)** |  |  |  |
| 5mC | Wild type | HC (5% CO<sub>2</sub>) | - |
|  | Wild type | LC (0.01% CO<sub>2</sub>) | - |
| **Histone modification (ChIP-Seq)** |  |  |  |
| H3K4me2 | Wild type | HC (5% CO<sub>2</sub>) | Abcam, ab7766 |
|  | Wild type | LC (0.01% CO<sub>2</sub>) | Abcam, ab7766 |
|  | Mutant M4 | LC (0.04% CO<sub>2</sub>) | CST, #9725 |
|  | Mutant M4 | Input (0.04% CO<sub>2</sub>) | - |
|  | Mutant M6 | LC (0.04% CO<sub>2</sub>) | CST, #9725 |
|  | Mutant M6 | Input (0.04% CO<sub>2</sub>) | - |
|  | Wild type | WT (0.04% CO<sub>2</sub>) | CST, #9725 |
|  | Wild type | Input (0.04% CO<sub>2</sub>) | - |
| H3K9ac | Wild type | HC (5% CO<sub>2</sub>) | Abcam, ab10812 |
|  | Wild type | LC (0.01% CO<sub>2</sub>) | Abcam, ab10812 |
| H3K27ac | Wild type | HC (5% CO<sub>2</sub>) | Millipore, 07-360 |
|  | Wild type | LC (0.01% CO<sub>2</sub>) | Millipore, 07-360 |
| Kcr | Wild type | HC (5% CO<sub>2</sub>) | PTM BioLab, PTM-501 |
|  | Wild type | LC (0.01% CO<sub>2</sub>) | PTM BioLab, PTM-501 |
| H3K9me3 | Wild type | HC (5% CO<sub>2</sub>) | Abcam, 8898 |
|  | Wild type | LC (0.01% CO<sub>2</sub>) | Abcam, 8898 |
| H3K27me3 | Wild type | HC (5% CO<sub>2</sub>) | Millipore, 07-449 |
|  | Wild type | LC (0.01% CO<sub>2</sub>) | Millipore, 07-449 |
| H2A.Z | Wild type | HC (5% CO<sub>2</sub>) | CST, #2718 |
|  | Wild type | LC (0.01% CO<sub>2</sub>) | CST, #2718 |
| Input1 | Wild type | HC (5% CO<sub>2</sub>) | - |
|  | Wild type | LC (0.01% CO<sub>2</sub>) | - |
| Input2 | Wild type | HC (5% CO<sub>2</sub>) | - |
|  | Wild type | LC (0.01% CO<sub>2</sub>) | - |
| **Nucleosome occupancy (MNase-Seq)** |  |  |  |
| Nucleosome | Wild type | HC (5% CO<sub>2</sub>) | - |
|  | Wild type | LC (0.01% CO<sub>2</sub>) | - |
| **Chromosome architecture (Hi-C)** |  |  |  |
| Spatial contact | Wild type | HC (5% CO<sub>2</sub>) | MboI |
|  | Wild type | LC (0.01% CO<sub>2</sub>) | MboI |

| **Transcriptome (RNA-Seq)** |  |  |  |
| --- | --- | --- | --- |
| mRNA | Mutant M4 | LC (0.04% CO<sub>2</sub>) | - |
|  | Mutant M6 | LC (0.04% CO<sub>2</sub>) | - |
|  | Wild type | LC (0.04% CO<sub>2</sub>) | - |

### Dissecting the Role of DNA Methylation in Low CO_2_ Adaptation

The DNA 5mC methylomes of *N. oceanica* under LC stress were examined first, as previous studies frequently suggest that DNA methylation is essential for gene expression regulation in almost all eukaryotes (Ferrari et al., 2023; Greenberg and Bourc’his, 2019; Steadman et al., 2020; Zhang et al., 2018). A high-resolution DNA methylation map that covers 91.6% of all cytosines in the genome was generated via whole-genome bisulfite sequencing (WGBS), which integrates six individual methylome profiles (**Table S1**). Surprisingly, only 0.13% of the cytosines in the genome were methylated (**Table S1**). Given that the conversion efficiency of bisulfite treatment in our samples was not perfect (Krueger et al., 2012; Zhou et al., 2020), the actual 5mC level in *N. oceanica* might still be overestimated. Further analysis revealed that, among the methylated cytosines in *N. oceanica*, approximately 51% were in the CHH context, whereas 29% and 20% were in the CHG and CpG contexts, respectively. This distinct level and distribution pattern of 5mC contrasts sharply with that observed in green algae such as *Chlamydomonas reinhardtii* and *Volvox carteri*, which possess relatively abundant 5mC (Babinger et al., 2001; Feng et al., 2010; Fu et al., 2015; Hattman et al., 1978). The extremely low abundance of 5mC in *N. oceanica* suggests a limited role for DNA methylation in the dramatic changes of gene expression during adaptation to LC conditions.

We next compared the levels of 5mC between HC and LC conditions and found no significant changes in different methylation contexts (CpG, CHG, and CHH), although an increased overall level of 5mC was observed (**Fig. S1A**). A detailed examination of the 5mC levels in various genomic regions, including genes, exons, protein-coding sequences (CDSs), transcription start sites (TSSs), transcription termination sites (TTSs), and transposon elements (TEs), also revealed limited changes (**Fig. S1B**). Additionally, no obvious correlation was found between 5mC occupancy and gene expression (**Fig. S1C**). Therefore, 5mC DNA methylation, exhibiting a limited regulatory effect on gene expression in *N. oceanica*, does not drive most LC-associated gene expression changes. This underscores the necessity of shifting our focus to other epigenomic layers to elucidate the regulatory mechanisms governing LC stress responses.

### Histone Modifications Orchestrate Transcriptional Regulation Under Low CO_2_

To elucidate the role of histone modifications in transcriptional regulation during CCM induction, we performed chromatin immunoprecipitation sequencing (ChIP-Seq) to map seven histone marks genome-wide under HC and LC conditions. Validation of antibody treatment efficacy via deepTools “fingerprints” analysis (Ramírez et al., 2016) confirmed enrichment of H3K4me2, H3K9ac, Kcr, and H3K27ac, which were then selected for further analysis (**Fig. S2A** and **Table S2**). Replicates exhibited strong correlations in terms of sequencing coverage (Pearson’s R > 0.93; **Fig. S2B**). Notably, these histone marks were enriched primarily in gene-rich regions (**Fig. S3A**) and were predominantly localized within promoter-TSS regions (60.6%-76.3%; **Fig. S3B-C**), indicating their critical functions in gene regulation and transcriptional initiation (Juven-Gershon and Kadonaga, 2010; Parvin et al., 1992).

Further review of the distribution patterns revealed significant enrichment of H3K9ac, H3K27ac, and Kcr at exactly TSSs (**Fig. 2A**), resembling patterns in land plants and metazoans (Prakash and Fournier, 2018; Zhou et al., 2011). In contrast, H3K4me2 displayed a distinct dual-peak profile around the TSS with relatively lower levels in close proximity to the TSS (**Fig. 2A**), resembling the observed patterns in animals (Hu et al., 2013; Liu et al., 2011; Rickels et al., 2016). Quantitative analysis revealed 5,587-6,180 peaks for H3K9ac, H3K27ac, and Kcr, whereas H3K4me2 exhibited 10,105 peaks due to its dual-peak pattern (**Table S3**). Intriguingly, our data revealed a high level of overlap among these histone marks, with significant co-occurrence at many gene loci (**Fig. S3D-E**). For instance, the NO20G00630 gene (NoCA5), a critical component of the *N. oceanica* CCM (Gee and Niyogi, 2017), exhibited overlapping marks of H3K9ac, H3K27ac, and Kcr, which were correlated with its significant upregulation (six-fold increase) under LC conditions (**Fig. 2B**).

**Figure 2.**
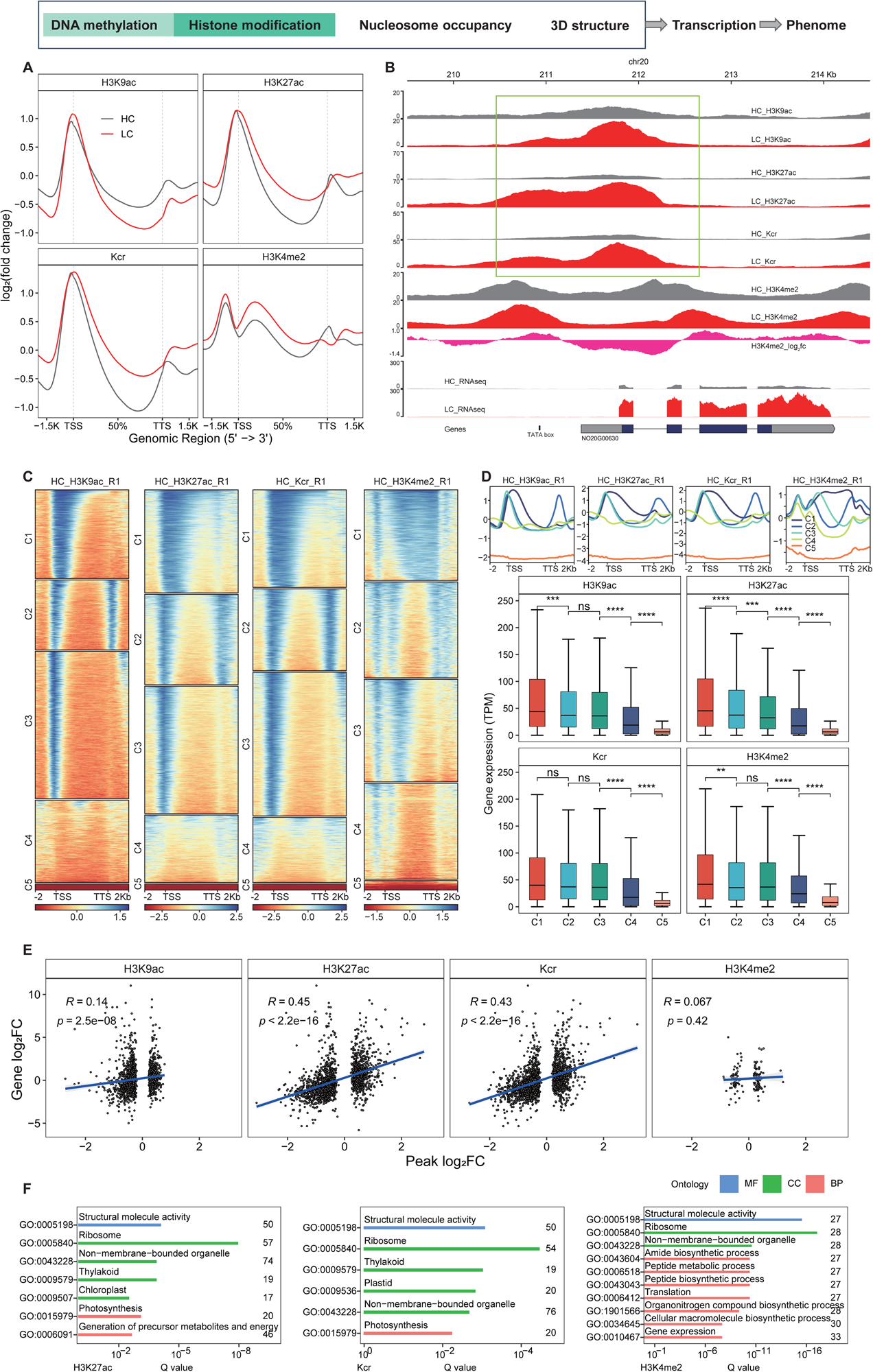
Dynamics of histone modification marks under HC and LC conditions in *N. oceanica*. **(A)** Enrichment profile of H3K4me2, H3K9ac, Kcr, and H3K27ac along genes under HC (upper) and LC (below) conditions. (**B**) Dynamics of histone modifications around the carbonate dehydratase gene NO20G00630 (NoCA5). Green box highlights the significantly changed peaks. **(C)** Genes were clustered into five groups via *k-*means algorithm based on distribution of H3K9ac, H3K27ac, Kcr, and H3K4me2 marks in HC and LC samples. Replicate 1 is shown. Genes were clustered based on HC samples. **(D)** Comparisons of gene expression for the five different clusters. **(E)** Scatter plots showing the relationships between changes in histone marks and gene expression. **(F)** Enrichment of GO terms associated with DHM-associated genes.

The co-occurrence of histone marks and their positive correlation with gene expression at the NoCA5 gene (**Fig. 2B**) prompted us to perform a genome-wide correlation analysis between gene expression and histone modifications. Indeed, genes marked by these modifications exhibited higher expression levels than those without such marks did (**Fig. S4A**). Genes were classified into five clusters (C1-C5) based on peak characteristics under HC conditions, with a decreasing order of histone mark abundance from C1 to C5 (**Fig. 2C**; **Fig. S4B**). Typically, the expression levels followed the order C1>C2≥C3>C4>C5, affirming the positive correlation between histone mark presence and gene expression (**Fig. 2D**; **Fig. S4C**). Overall, higher expression was noted in the C1-C3 clusters enriched with histone marks, with C5 genes, mostly chloroplast and mitochondrial genes, exhibiting the lowest expression and a lack of histone marks.

Given the observed positive correlation between histone marks and gene expression, we proceeded to investigate differential histone modifications between HC and LC conditions to better understand transcriptional reprogramming. We identified 1,823-2,452 differential peaks (differential histone marks, DHMs) for H3K9ac, H3K27ac, and Kcr, but only 263 for H3K4me2 (**Table S4**). Linear regression of DHM peak abundance changes against gene expression changes revealed positive correlations for all histone marks, with a noticeably lower correlation for H3K4me2, likely due to the fewer number of DHMs identified for this modification (**Fig. 2E**). Functional annotation of DHM-associated genes (DHMGs) revealed enrichment in molecular functions and cellular components related to ribosomal biogenesis and organelle function. DHMGs enriched in H3K27ac and Kcr were linked to photosynthesis processes, whereas those enriched in H3K4me2 were correlated with various biological processes, including gene expression, translation, biosynthesis of peptide, organonitrogen and cellular macromolecule, etc. (**Fig. 2F**). Collectively, our findings underscore the prominent roles of histone modifications in orchestrating gene expression reprogramming in *N. oceanica* under LC stress, offering insights into the underlying epigenetic mechanisms involved.

### Nucleosome Dynamics Underlie Altered Gene Expression in Response to Low CO_2_

The distinct roles of two different epigenetic layers, with limited effects on 5mC DNA methylation and prominent regulatory roles of histone modifications, revealed a unique epigenomic regulation system in *N. oceanica* that differs from that of other microalgal models, such as *Chlamydomonas reinhardtii*. Thus, we next asked whether nucleosome positioning, another critical factor regulating gene expression, might play a role in adapting to LC conditions via micrococcal nuclease sequencing (MNase-Seq) experiments under HC and LC conditions. High reproducibility among biological replicates was achieved, as evidenced by a Pearson coefficient of 0.93 for genome-wide read coverage (**Fig. S5A**). We identified 81,031 nucleosomes under HC and 87,845 under LC (**Table S5**; **Fig. S5B**), with the nucleosome repeat length (NRL) peaking consistently at 178 bp under both conditions, indicating stability in the nucleosome arrangement (**Fig. 3A**). Interestingly, regions adjacent to the TSS and TTS showed significant nucleosome depletion, pinpointing these areas as critical for transcriptional regulation (**Fig. 3B**).

**Figure 3.**
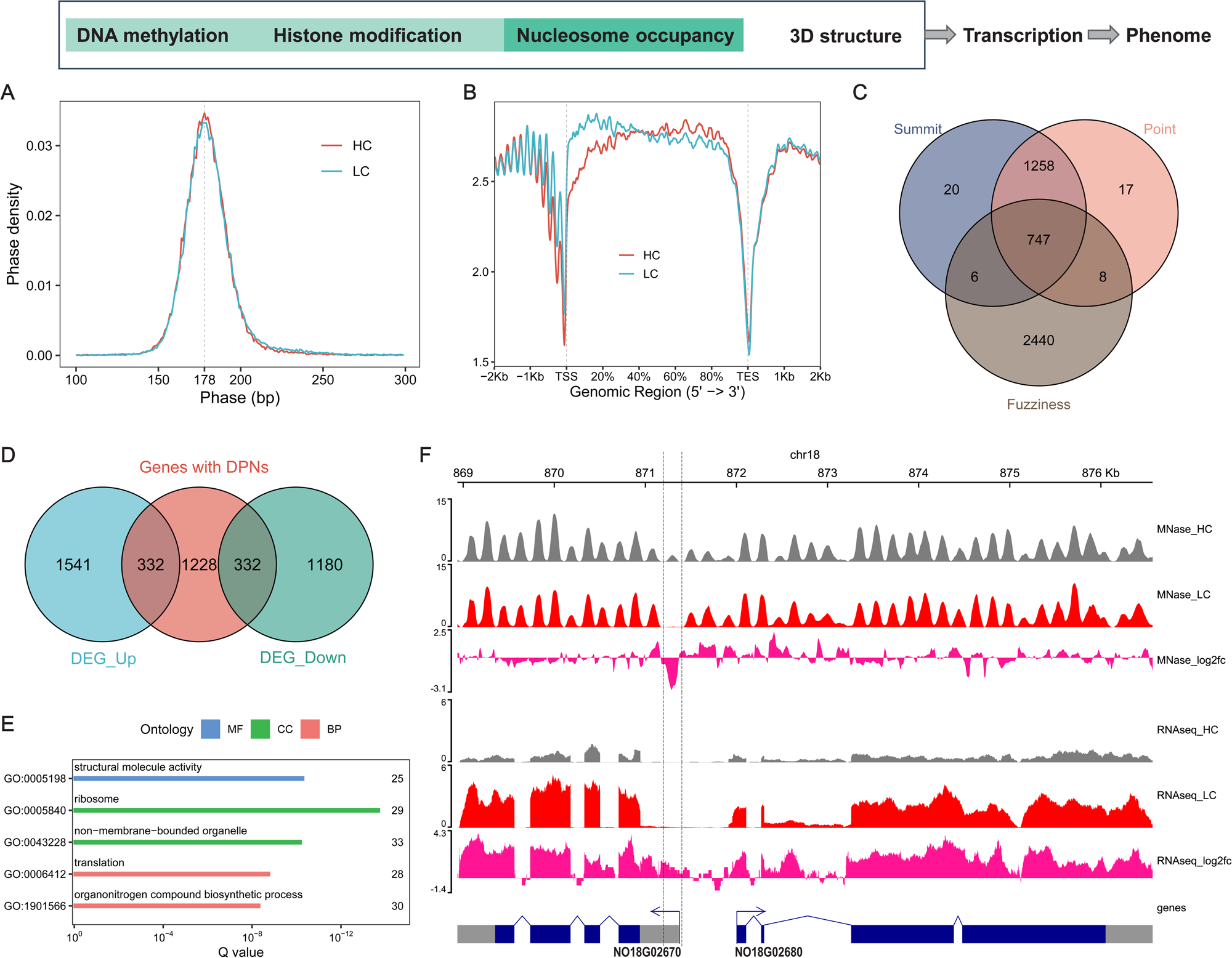
Dynamics of nucleosome occupancy under HC and LC conditions in *N. oceanica*. (**A**) Nucleosome phases peaked at 178 bp under both HC and LC. (**B**) Nucleosome distribution along genes under HC and LC conditions. (**C**) Venn diagram illustrating the overlap between three categories of differentially positioned nucleosomes (DPNs). (**D**) Correlating the DPNs with the DEGs. DPNs were mapped to TSS regions of genes (1 kb upstream to TSS). (**E**) Enrichment of GO terms in downregulated genes associated with dynamic nucleosomes in promoter regions. (**F**) The loss of a nucleosome results in the upregulation of the adjacent gene pair NO18G02670 and NO18G02680, which is promoter-overlapping. NO18G02680 is a Zn_2_Cys_6_ fungal-type transcription factor.

A comparison between LC and HC revealed 4,496 differentially positioned nucleosome (DPN) peaks under LC stress (FDR < 0.01; **Fig. 3C**), with the majority located in gene promoters. These DPNs were mapped to 1,892 genes, of which a significant proportion (35.1%) were differentially expressed (**Supplemental file 1**). The enrichment of DEGs among those containing DPNs (Fisher’s exact test, *p* < 0.01) underscores the pivotal role of nucleosome positioning in gene regulation during LC adaptation in *N. oceanica*. Further analysis demonstrated a balanced distribution of upregulated (17.5% versus 17.8% genome-wide; hypergeometric test, *p* = 0.77) and downregulated genes (17.5% versus 14.6% genome-wide; hypergeometric test, *p* < 0.01) (**Fig. 3D**). Gene Ontology (GO) analysis of these genes revealed enrichment of ribosomal components among the downregulated genes, suggesting broad reprogramming toward reduced metabolic activity in response to LC stress (**Fig. 3E**; **Table S6**). Notably, the upregulation of the Zn_2_Cys_6_ fungal-type transcription factor (TF) NO18G02680, which is homologous to Naga_100262g2 in *N. gaditana* and may act as a regulator of LC stress adaptation, was apparent (**Fig. 3F**). In *N. gaditana*, another Zn_2_Cys_6_ TF (Naga_100104g18; homolog to NO09G01030 in *N. oceanica* IMET1) regulates lipid metabolism without affecting growth (Ajjawi et al., 2017). Additionally, DPNs were associated with a bicarbonate transporter (NoBCT2, NO01G02900), which is critical for the biophysical CCM. Moreover, genes such as argininosuccinate synthase (ASS, NO05G00910) and ornithine decarboxylase (ODC, NO05G00760) in the ornithine urea cycle (OUC) also showed associations with DPNs, suggesting their involvement in the adaptive response to LC conditions.

### Chromatin State Alternation in Response to Low CO_2_ Adaption

Since both histone modifications and nucleosome positioning significantly affect gene expression regulation (**Fig 2** and **Fig 3**), we then integrated these two data types and examined the changes in the chromatin state (CS) under LC stress. Specifically, five CSs were identified, each representing a distinct combination of nucleosome occupancy and histone marks (**Fig 4A**). CS1 was deficient in all four histone marks, whereas CS2 exhibited higher nucleosome occupancy than did CS1. CS3 was primarily marked by H3K4me2, CS4 was enriched in all histone modifications, while CS5 was enriched in H3K9ac, H3K27ac, and Kcr (**Fig. 4A**). The separate distribution pattern of H3K4me2 as represented by CS3 suggests that H3K4me2 does not always overlap with the other three histone marks and shows a relatively distinct distribution pattern in some genomic regions. In addition, these states exhibited distinct positional preferences across the genome under both HC and LC conditions (**Fig. 4B**). For instance, CS1 was the predominant state, encompassing 36% and 42% of the *N. oceanica* genome under HC and LC conditions, respectively. In contrast, CS5, representing approx. 10% of the genome, was highly concentrated at the TSS.

**Figure 4.**
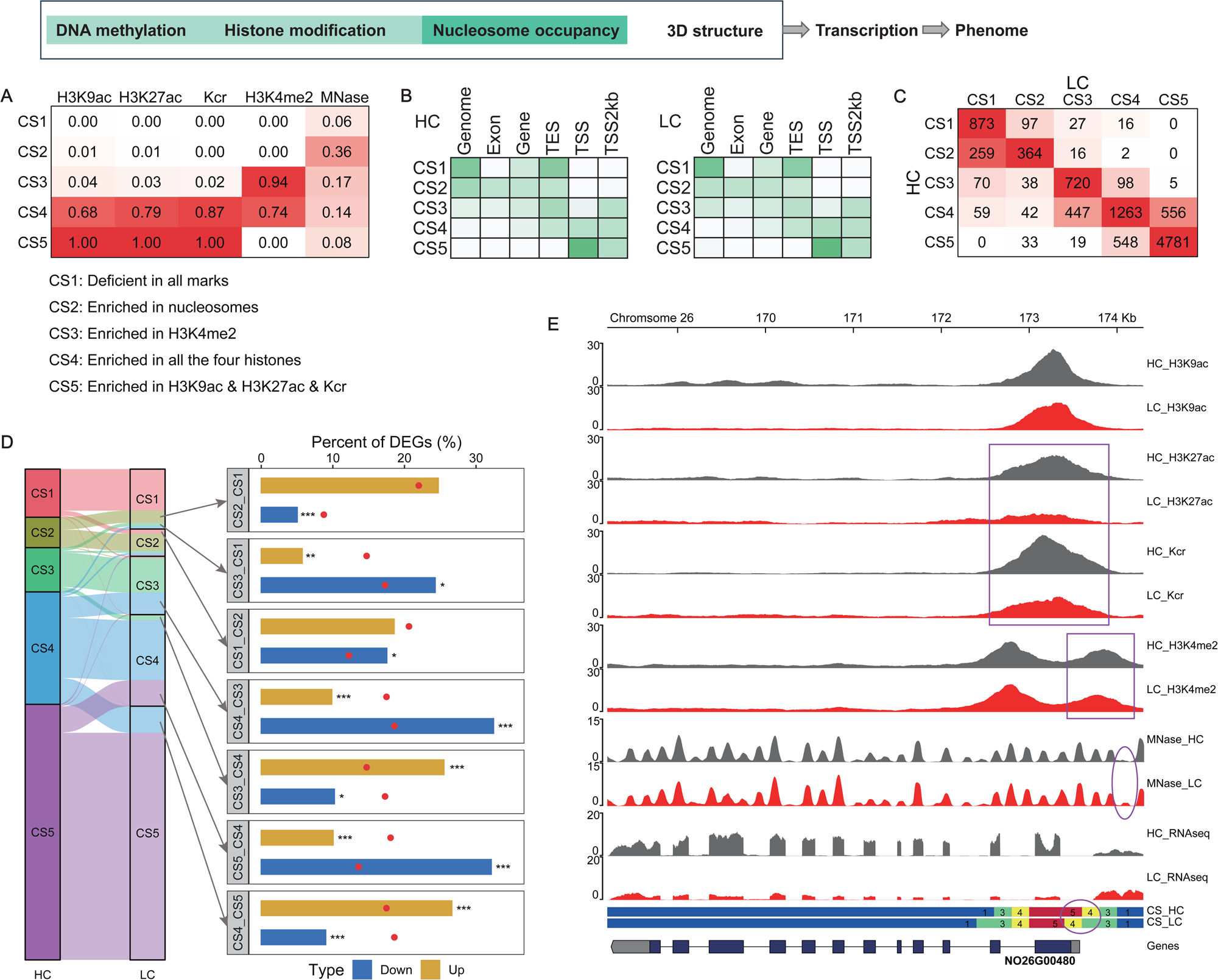
Dynamics of chromatin states under HC and LC conditions in *N. oceanica*. (**A**) The emission parameters derived from the Hidden Markov Model constructed through chromHMM analysis represent the characteristics of the five distinct chromatin states (CSs). For the HC and LC samples, the CS of each genomic fragment was defined by the peak occurrence of H3K4me2, H3K9ac, Kcr, H3K27ac, and nucleosome occupancy. The darker red color corresponds to a greater probability of observing the epigenetic mark in the state. (**B**) Overlap fold enrichment of various genomic regions with five chromatin states in the HC and LC samples. Relative enrichment value was used, which subtracts the minimum value in the column and then divides by the maximum column value. White corresponds to 0, and the darkest value is the maximum value after transformation anywhere in the heatmap. (**C**) Transition matrix based on the chromatin states of genes (defined as the chromatin state of TSS). (**D**) Percent of DEGs showing changes in CS. Genes were binned according to the CS of the HC samples, with *p*-values calculated using the perturbation test. (**E**) Dynamics of epigenetic signals and chromatin states around the malic enzyme-encoding gene NO26G00480.

Furthermore, our integrated analysis revealed the dynamic nature of CS alterations during the transition from HC to LC conditions. A notable proportion of genes exhibited CS shifts, especially at the TSS, indicating active epigenetic reprogramming (**Fig. 4C**). Approximately 26.8% of the DEGs underwent CS changes, underscoring their epigenetic regulation during LC adaptation. The transitions from CS4 to CS1, CS4 to CS3, and CS5 to CS2 were associated with significant gene downregulation, likely due to the loss of active histone modifications near the TSS (**Fig. 4D**). Conversely, transitions between CS4 and CS5 were linked to both gene upregulation and downregulation, indicating a nuanced role in transcriptional regulation (**Fig. 4D**).

GO enrichment analysis further highlighted the functional implications of these CS changes. For example, genes undergoing transitions from CS5 to CS4 were enriched for ribosomal components, such as those involved in the ribosome (GO:0005840), implying a broad reprogramming toward reduced metabolic activity in response to LC stress. This finding was corroborated by transcriptomic data, which showed consistent downregulation of these ribosomal genes under LC (**Table S7**). Additionally, key genes such as NoCA5, which is essential for the CCM, and others involved in the ornithine urea cycle (e.g., ornithine decarboxylase, ODC) exhibited CS shifts, further emphasizing the complex interplay between chromatin dynamics and gene regulation under varying CO_2_ conditions.

Specifically, the malic enzyme 1 (ME1) gene, which catalyzes the oxidation of malate to pyruvate with CO_2_ release, showed remarkable CS changes under LC (**Fig. 4E**). This gene was downregulated, which correlated with decreased levels of H3K27ac and Kcr. Moreover, an H3K4me2 peak for ME1 was repositioned closer to the TSS, and the adjacent nucleosome became more inaccessible (**Fig. 4E**), suggesting multiple transcriptional regulatory pathways. Thus, the multi-faceted analyses of *N. oceanica* not only reveal immediate CSs in response to LC but also provide comprehensive insights into the epigenetic landscapes that underpin adaptive gene regulation.

### Reorganization of 3D Chromosome Architecture in Response to Low CO_2_ Adaption

On top of the fine-level CS changes that encompass 5mC DNA methylation, histone modifications, and nucleosome positioning, we further defined the role of higher-order 3D chromatin conformation in this LC adaption process by generating high-resolution Hi-C interaction maps via Hi-C sequencing of *N. oceanica* under HC and LC conditions (**Fig. 5A**; **Table S8** and **Fig. S6A-B**; **Methods**). The 3D genome architecture (**Fig. 5B**) revealed that each chromosome folds in half, with telomeres colocalizing and centromeric regions displaying fusion, which resembles the lattice structure of haploid yeast (Shao et al., 2018) and suggests susceptibility to large-scale chromosomal manipulations. Consistent with eukaryotic chromatin organization (Lieberman-Aiden et al., 2009), *N. oceanica* chromosomes can be divided into A/B compartments (**Fig. 5C**, top panel), where interactions occur predominantly within the same compartment. Compartment A aligns with more central chromosomal regions, which are associated with open chromatin and higher gene expression, whereas compartment B is located near chromosomal tails, denoting closed chromatin (Wilcoxon test, *p* < 0.0001; **Fig. 5C**, right bottom panel).

**Figure 5.**
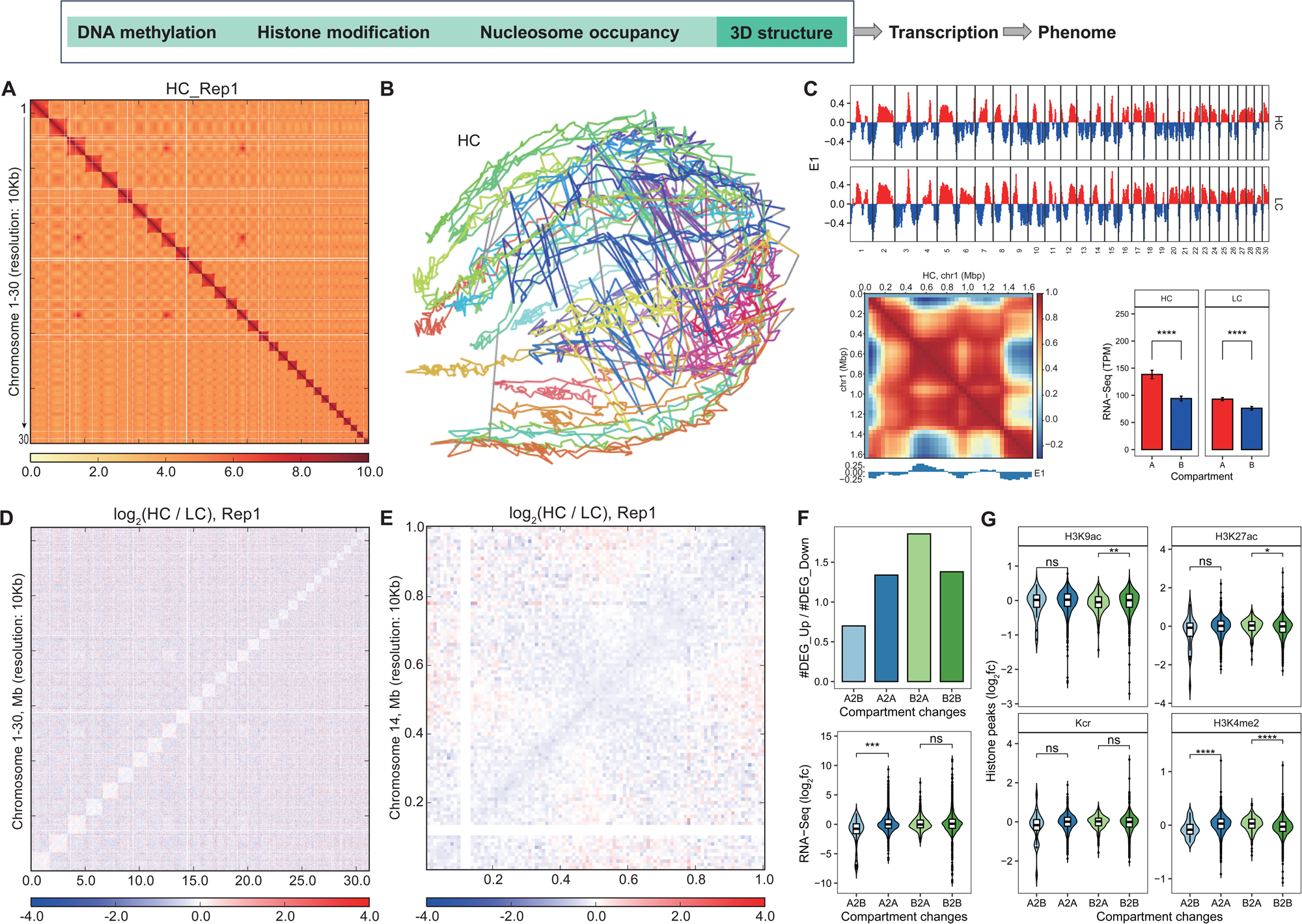
Dynamics of 3D chromatin architecture under HC and LC conditions in *N. oceanica*. **(A)** The interaction map of *N. oceanica* at 10 kb resolution. (**B**) 3D structure of the *N. oceanica* genome reconstructed from the normalized genome-wide interaction matrix. Different chromosomes are shown in different colors. (**C**) Chromosome-specific A/B compartment profiles of *N. oceanica* affect gene transcription under HC and LC conditions (at 40 kb resolution). Top: compartment A/B segmentations across the *N. oceanica* genome; Left bottom: Pearson correlation matrix of observed/expected Hi-C interactions and the resulting compartment segmentation for chromosome 1 and under HC conditions; Right bottom: expression comparisons between genes in different compartments for HC and LC conditions. (**D**) The difference interaction matrix for all chromosomes at 10 kb resolution between HC and LC conditions for Replicate 1. (**E**) The difference interaction matrix for chromosome 14 at 10 kb resolution between HC and LC conditions for replicate 1. (**F**) Dynamics of local compartments in *N. oceanica* correlate with differential expression of genes. (**G**) Dynamics of local compartments in *N. oceanica* correlate with histone modification changes.

A comparison of the HC and LC contact maps at 10 kb resolution revealed that although the higher-order chromatin architecture was conserved (**Fig. 5D**, **Fig. S6C-D**), extensive local chromatin architecture dynamics did occur (**Fig. 5E**). Intriguingly, such dynamics aligned with gene expression alterations, where A to B compartment transitions were associated with a higher incidence of downregulated genes and a notable decrease in overall expression (“A2B” vs “A2A”; Wilcoxon test, *p* < 0.001). In contrast, B to A transitions were correlated with an increase in upregulated genes, albeit without significant changes in expression levels (“B2A” vs “B2B”; Wilcoxon test, *p* > 0.05) (**Fig. 5F**). In addition, among all the histone marks examined, chromosomal compartment shifts were revealed by H3K4me2 distribution patterns (**Fig. 5G**). Specifically, a significant reduction in H3K4me2 alterations was observed in the A2B transition group compared with the A2A group (Wilcoxon test, *p* < 0.0001), and a marked increase was noted in the B2A transition group relative to the B2B group (Wilcoxon test, *p* < 0.0001). As H3K4me2 is also recognized as an active enhancer mark (Chepelev et al., 2012; Ong and Corces, 2012; Pekowska et al., 2011; Wang et al., 2008), we investigated whether distal regulation through spatial contact loops between promoters and enhancers also contributes to gene regulation (Grubert et al., 2020; Hamamoto and Fukaya, 2022; Zagirova et al., 2024). Several significant “mid-range” chromatin contacts were identified from the HC (10 contacts) and LC (24 contacts) conditions (4∼835 kb, FDR < 0.05), with 11 out of the 21 genes involved in these contacts (52.4%) being significantly regulated. Furthermore, these chromatin contacts were associated with the promoter regions of 10 genes (**Fig. S6E**), indicating the implication of enhancer/promoter-promoter interactions in response to LC adaption in *N. oceanica*. These findings suggest that changes in histone modifications (especially H3K4me2) may underpin compartment transitions, thereby influencing gene expression dynamics under LC stress.

### Functional Validation of Epigenetic Regulators in Carbon Assimilation

Given the intricate epigenetic landscape unraveled by the comprehensive profiling, we next investigated the carbon assimilation mechanisms in *N. oceanica* by pinpointing a specific histone mark. H3K4me2 has emerged as a prime candidate: (*i*) H3K4me2 has a distinctive distribution pattern prominently observed in the genomic regions of active CSs (**Fig. 2A** and **Fig. 4A**); (*ii*) H3K4me2 is a key epigenetic marker in our 3D chromatin compartmental analysis, linked to spatial genome organization and gene expression dynamics (**Fig. 5G**); and (*iii*) H3K4me2 has a unique dual-peak profile around the TSS (**Fig. 2A**), which is distinct from that of other microalgae or plants. These findings suggest a unique and complex role of H3K4me2 in the transcriptional regulation of *N. oceanica*.

To validate the function of H3K4me2 in *N. oceanica*, a candidate H3K4me2 methyltransferase (NO24G02310) was knocked out via CRISPR/Cas9, resulting in two frameshift mutants (M4 and M6) with protein truncation before the functional domains (**Fig. 6A**). H3K4me2 ChIP-seq profiling revealed largely similar yet distinct distribution patterns between the mutants and WT, with numerous differential peaks observed (**Fig. S7A-C** and **Fig. 6B**). Notably, H3K4me2 peaks in the mutants shifted closer to the TSS on a genome-wide scale, indicating that the candidate methyltransferase significantly influences H3K4me2 positioning (**Fig. S7A** and **Fig. S7D**). Under LC, M4 and M6 exhibited significant reductions in both growth (20.0% and 21.6%, respectively) and biomass productivity (14.7% and 10.1%, respectively), along with decreased Fv/Fm (5.7% and 11.3%, respectively) (**Fig. 6C-E**). However, these phenotypic changes disappeared under HC conditions, with no differences in growth, biomass, or photosynthetic efficiency compared with those of the WT, except for a slight increase in the Fv/Fm of the M4 mutant at 2-5 days and a 2.6% reduction in M6 (**Fig. 6F-H**). These results suggest that H3K4me2 methyltransferase knockout impairs carbon assimilation and highlight the essential role of H3K4me2 in LC adaptation in *N. oceanica*.

**Figure 6.**
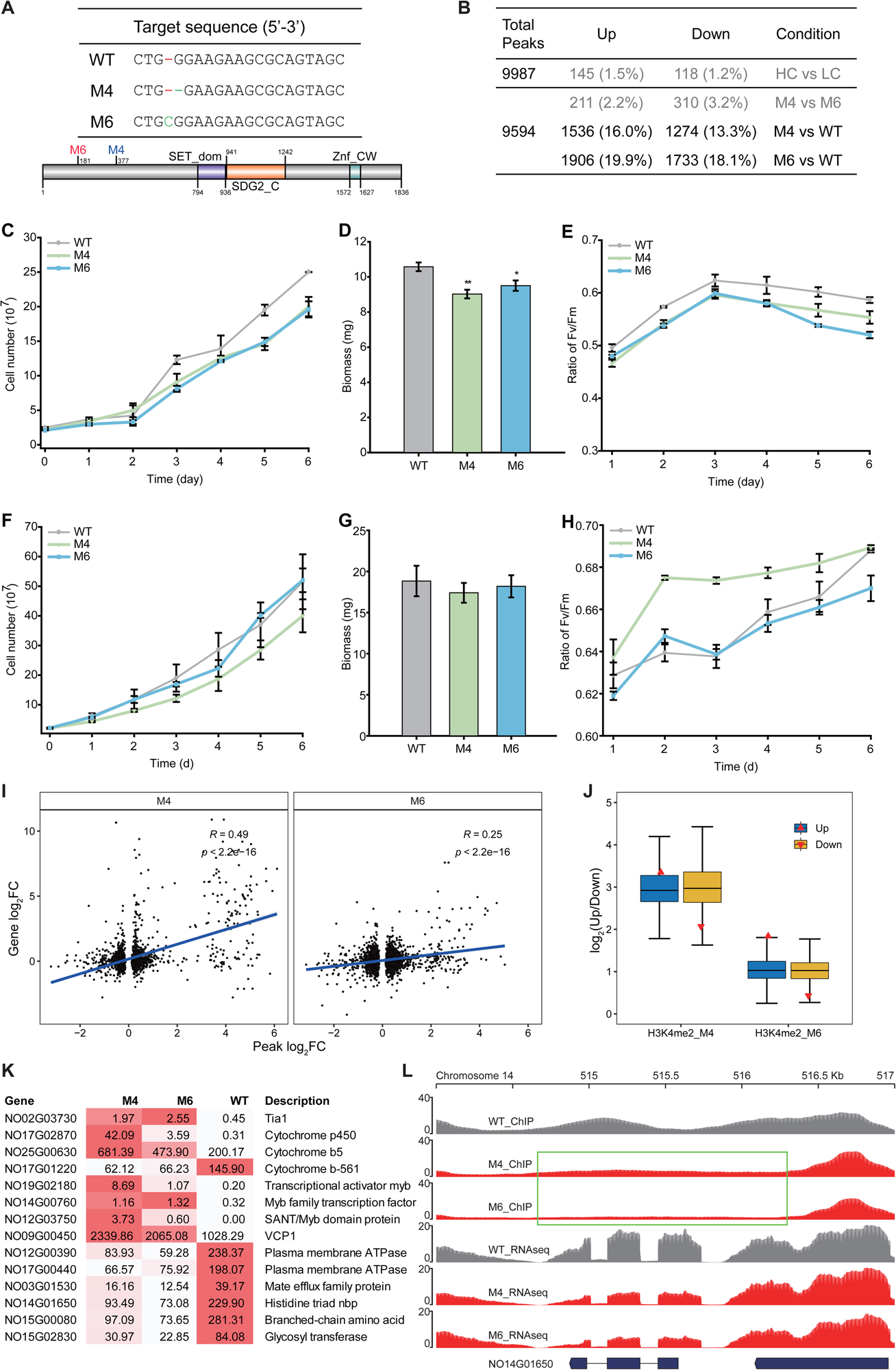
NO24G02310 knockout revealed that H3K4me2 mediates carbon assimilation. (**A**) The genome sequence of NO24G02310-knockout mutants at the gRNA target sites. The mutants were truncated before the functional domains of NO24G02310. (**B**) Based on the number of significantly changed peaks, the gene knockout caused a higher degree of dynamics of H3K4me2 marks than CCM induction did. (**C-E**) The growth phenotype, biomass productivity, and photosynthesis parameters (Fv/Fm) of the mutants under low CO_2_ (0.04%). (**F-H**) The corresponding phenotypes under high CO_2_ (5%). (**I**) Links between the dynamics of histone marks and gene expression. (**J**) Association between differential gene expression and peak shift of H3K4me2 marks. (**K**) Expression profiles of selected genes that potentially contribute to the phenotype of NO24G02310-knockout mutants. The average TPM values are shown. (**L**) Dynamics of H3K4me2 histone modifications around the NoHINT (NO14G01650) gene. Green box: the region of disappeared peaks.

Transcriptomic analyses of both the mutants and the WT identified 448 DEGs (M4 vs. WT) and 423 DEGs (M6 vs. WT), clearly distinguishing the mutants from the WT (**Fig. S7E-F**). Linear regression analysis revealed a positive correlation between changes in H3K4me2 peak abundance and gene expression, confirming the role of H3K4me2 in gene activation (*p* < 0.01; **Fig. 6I**). Furthermore, both the upregulated and downregulated peak groups had significantly higher ratios of DEGs than random sampling groups, reinforcing the impact of H3K4me2 on gene expression (*p* < 0.01; **Fig. 6J**). The transcriptomic profiles under LC revealed intriguing trends among the DEGs (**Fig. 6K**). For example, TIA1, which is a protein involved in stress granule formation and stress response, was upregulated, suggesting a highly stressed state that characterizes the mutants. Additionally, the histidine triad nucleotide-binding protein NoHINT (NO14G01650) exhibited an absence of H3K4me2 peaks and significantly reduced expression (log_2_FC = -1.49), indicating its critical role in the growth and carbon assimilation of the mutants (**Fig. 6L**). An examination of published transcriptome datasets deposited in the NanDeSyn database (Gong et al., 2020) revealed that NoHINT was specifically LC-downregulated (log_2_FC = -1.29) and remained unaffected by other stress conditions. This unique response underscores the potential role of NoHINT in H3K4me2-mediated regulation of carbon assimilation in *N. oceanica*.

Concurrently, two plasma membrane H^+^-ATPases (PMAs), which are potentially crucial for nutrient uptake, intracellular pH regulation and CCMs, were downregulated in the mutants. Intriguingly, while NoPMA1 expression remained stable across HC and LC conditions, NoPMA2 was consistently downregulated (LC vs HC, log_2_FC = -1.35), mirroring observations in *C. reinhardtii* where PMA under-regulation was associated with impaired adaptation to high CO_2_ levels (Choi et al., 2021).

The convergence of these expression patterns, coupled with the absence of H3K4me2 peaks at the NoHINT locus, strongly implicates both NoHINT and NoPMA2 as key targets of H3K4me2 methylase, potentially exerting a significant influence on LC adaptation in *N. oceanica*. To verify these functional roles of NoHINT, we generated two knockout mutants, NoHINTko-1 and NoHINTko-2 (**Fig. 7A**). Under LC, these mutants exhibited a subtle yet significant growth reduction (5.4-14.0%, *p* < 0.05) between days 3-7 of cultivation compared to WT, despite no substantial differences in final biomass or Fv/Fm after nine days (**Fig. 7B-D**). This transient growth impairment, particularly evident during nutrient-replete conditions, supports the putative role of NoHINT in modulating carbon assimilation, possibly through specific interactions with growth-related enzymes.

**Figure 7.**
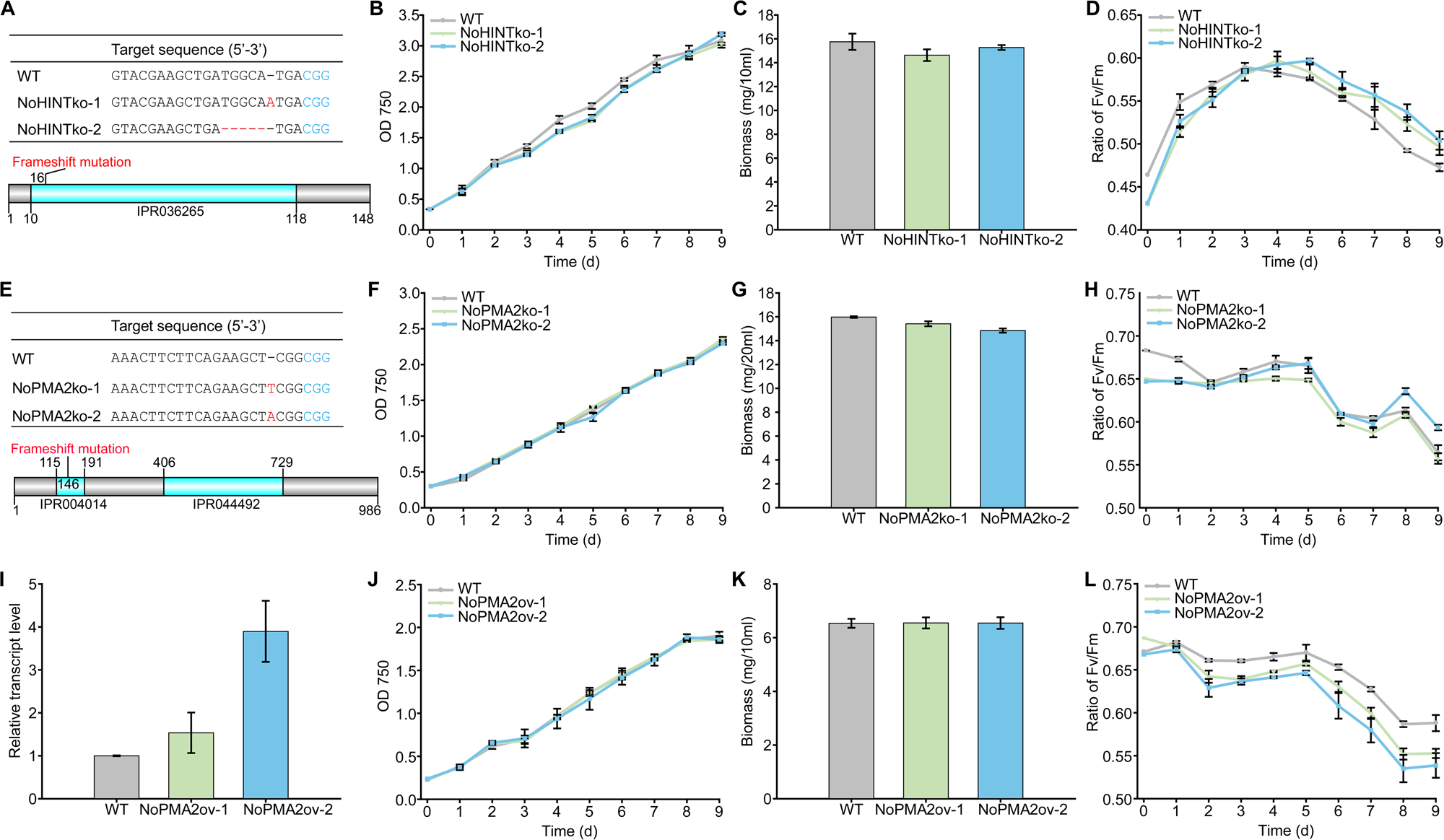
Knockout and overexpression mutants of NoHINT and NoPMA2 reveal epigenetics-mediated regulatory mechanisms of carbon assimilation in *N. oceanica*. (**A**) qPCR analysis of NoHINT overexpression strains. (**B-D**) Evaluation of the growth phenotype, biomass productivity, and photosynthesis parameters (Fv/Fm) of NoHINT overexpression strains. (**E**) Genotype analysis of NoPMA2 knockout mutants. (**F-H**) Assessment of the growth phenotype, biomass productivity, and photosynthesis parameters (Fv/Fm) of NoPMA2 knockout mutants. (**I**) qPCR analysis of NoPMA2 overexpression strains. (**J-L**) Evaluation of the growth phenotype, biomass productivity, and photosynthesis parameters (Fv/Fm) of NoPMA2 overexpression strains.

For NoPMA2, both knockout (NoPMA2ko-1 and NoPMA2ko-2; **Fig. 7E**) and overexpression (NoPMA2ov-1 and NoPMA2ov-2; **Fig. 7I**) mutants were generated. Under LC, while knockout mutants showed no significant alterations in growth or biomass, a marginal increase in Fv/Fm (5.2%) was observed in NoPMA2ko-2 (**Fig. 7F-H**). Conversely, NoPMA2 overexpression strains presented no differences in growth or biomass but decreased Fv/Fm values (6.0% and 8.4% for NoPMA2ov-1 and NoPMA2ov-2, respectively; **Fig. 7J-L**). These photosynthetic efficiencies suggest that NoPMA2 may regulate photosynthesis by modulating pH gradients across chloroplast membranes.

Collectively, our findings provide compelling evidence that H3K4me2 regulates carbon assimilation in *N. oceanica* by targeting NoHINT and NoPMA2. The subtle yet consistent phenotypic alterations observed in the mutant strains underscore the fine-tuned nature of epigenetic regulation in microalgal carbon metabolism, and paves the way for targeted epigenetic modifications to increase microalgal productivity.

## Discussion

The epigenome, characterized by hierarchical layers of regulation from fine-scale modifications such as DNA methylation (Luo et al., 2015), histone modifications, and nucleosome dynamics to complex higher-order chromatin conformations, plays a pivotal role in gene expression regulation (Casas-Mollano et al., 2008; Kouzarides, 2007; Xiong et al., 2016; Zhang et al., 2009). However, for *Nannochloropsis* spp. which offer great potential for industrial CO_2_-to-oil conversion (Radakovits et al., 2012) and occupy a unique position in the Heterokonts by lacking chlorophyll *c* (unlike diatoms and close relatives; (Koh et al., 2019)), the complete epigenetic landscape remains largely unexplored. Here, we comprehensively characterized the structural and chemical aspects of epigenomics in *N. oceanica*, focusing on CCM, and analyzed them along with genomic and transcriptomic data following a shift to LC conditions. Our data revealed epigenomic profiles of 3D chromatin landscapes, nucleosome occupancy, DNA methylomes, and histone modifications including acetylation, methylation, and crotonylation, which shed light on epigenetic mechanisms associated with carbon assimilation, focusing on CCM. Moreover, through genetic manipulation of a candidate H3K4me2 methyltransferase and multi-omics analysis, our study uncovered the critical role of H3K4me2 in the regulation of carbon assimilation by targeting key genes, including NoHINT and NoPMA2. Functional validation of these identified targets via targeted gene knockout and overexpression revealed their subtle yet notable effects on growth and photosynthetic efficiency. Therefore, we have identified distinctive signatures and highlighted the active involvement of epigenetic regulation in *Nannochloropsis* under LC stress.

*Nannochloropsis oceanica* exhibits unique epigenetic signatures compared to higher plants and other algae. Specifically, H3K4me2 peaks show notable enrichment in promoters and gene body regions of active or poised genes, in contrast with the enrichment at proximal promoters and transcription start sites in higher plants (Liu et al., 2019). This dual-peak enrichment pattern aligns more closely with that observed in animals (Hu et al., 2013; Rickels et al., 2016) and the brown alga *Ectocarpus* (Bourdareau et al., 2021), yet diverges significantly from *Chlamydomonas reinhardtii* and plant patterns (Liu et al., 2019; Ngan et al., 2015). H3K4 methylation, which is generally facilitated by SET domain-containing methyltransferases, has diverse functional roles due to its differential evolution. Notably, the H3K4me2 methyltransferase NO24G02310 in *N. oceanica* features a SET domain (IPR001214), a zinc-finger domain (IPR011124), and a chromo-like domain (IPR016197). Functionally, H3K4me2 in *Nannochloropsis* promotes gene transcription, akin to its role in animals. In contrast, its function in plants remains debated, with studies in rice showing an inverse correlation with gene transcription during development (Liu et al., 2019), and in cotton (*Gossypium hirsutum*) it plays a role in the heat stress response (He et al., 2022).

In addition, *N. oceanica* displays an exceptionally low level of genomic 5-methylcytosine (∼0.1%), markedly lower than the methylation levels observed in animals, plants, and even closely related diatoms such as *Phaeodactylum tricornutum*, where methylated regions account for approximately 5% of global methylation (Fan et al., 2020; Feng et al., 2010; Lopez et al., 2015; Veluchamy et al., 2013). This deficiency is likely attributed to the absence of key DNA methyltransferases (DNMTs) such as DNMT2 and DNMT3, with only a candidate DNMT1 (NO12G03180) identified, paralleling the low methylation landscape observed in *Saccharomyces* (He et al., 2011). The near absence of methylation, potentially exacerbated by imperfect bisulfite conversion, underscores a unique epigenetic architecture that distinguishes *N. oceanica* from both green algae (such as *Chlamydomonas*) and diatoms, emphasizing its distinctive genomic regulation.

In the higher level of chromatin structures, we propose that histone marks of H3K4me2 are associated with the spatial genome organization of *N. oceanica*, especially in terms of compartment switching. Past studies highlighted the role of H3K9 methylation in chromatin compartmentalization in animals (Bian et al., 2020; Wang et al., 2019), with H3K27ac and H3K36me3 emerging as critical for A/B compartment prediction (Zheng et al., 2024). Our findings underscore the complex interplay between histone modifications and chromatin architecture in *Nannochloropsis*, necessitating further high-resolution studies. At the fine scale of chromatin structure, *N. oceanica* notably features a nucleosome repeat length of 179 bp, with open nucleosomes facilitating gene transcription at the promoter and TTS regions (**Fig. 3A-B**). In diatoms such as *Phaeodactylum tricornutum*, nucleosome depletion primarily occurs around 150 bp upstream of the TSS in highly expressed genes, but not at the TTS (Veluchamy et al., 2015). In plants, nucleosome positioning critically impacts gene expression, evolution, and alternative splicing (Jabre et al., 2021; Zhang et al., 2015). Remarkably, nucleosome depletion around the TTS is rare, observed notably in *Nannochloropsis* spp. and maize (Chen et al., 2017a), suggesting unique regulatory mechanisms in *N. oceanica*.

Based on this comprehensive epigenomic map, we analyzed the epigenetic patterns of key CCM components and assessed their epigenetic contributions to LC adaption (**Table S9**). It was remarkable that most of the key CCM genes that were differentially expressed presented at least one type of epigenetic regulation (**Fig. S8**). These epigenetic signatures play critical roles in determining gene transcription efficiency and orchestrating carbon assimilation traits in *N. oceanica*. Amid the crucial yet challenging task of establishing a causal relationship between phenotype alterations and specific chromatin-modifying enzymes, this study demonstrates the substantial potential of epigenetic breeding by confirming the influence of a candidate H3K4me2 methyltransferase on the carbon assimilation phenotype of *N. oceanica*. Additionally, based on genetic manipulation of machineries that underlie H3K4me2, we proposed a comprehensive model for the epigenetic regulation of CCM in *N. oceanica*, where NoHINT regulates growth and NoPMA2 influences photosynthesis. Thus, to enhance carbon fixation in *Nannochloropsis*, CCM is an important target for (epi)genetic engineering, and the epigenetic regulation could provide important clues for engineering schemes.

Taken together, this study provides the first panoramic epigenomic map of carbon assimilation under LC stress in photosynthetic organisms, and establishes a model for the epigenetic regulation of CCM in *N. oceanica*. These findings pave the way for enhancing carbon fixation and biomass productivity through targeted epigenetic modifications of industrial microalgae.

## Materials and Methods

### Algae cultivation

*Nannochloropsis oceanica* IMET1 was inoculated into the modified f/2 liquid medium, which was prepared with 35 g L^-1^ sea salt (Realocean, USA), 1 g L^-1^ NaNO_3_, 67 mg L^-1^ NaH_2_PO_4_*H_2_O, 3.65 mg L^-1^ FeCl_3_*6H_2_O, 4.37 mg l^-1^ Na_2_EDTA*2H_2_O, a trace metal mixture (0.0196 mg L^-1^ CuSO_4_*5H_2_O, 0.0126 mg L^-1^ NaMoO_4_*2H_2_O, 0.044 mg L^-1^ ZnSO_4_*7H_2_O, 0.01mg L^-1^ CoCl_2_, and 0.36 mg L^-1^ MnCl_2_*4H_2_O), and a vitamin mixture (2.5 µg L^-1^ VB_12_, 2.5 µg L^-1^ biotin, and 0.5 µg L^-1^ thiamine HCl) (Kang et al., 2015). The cells were first cultured in f/2 medium at 25 °C with 80±5 μmol m^-2^ s^-1^ continuous irradiation in a 1 L column reactor (inner diameter: 5 cm). The seed cultures were bubbled with 5% CO_2_. At the logarithmic growth phase (OD_750_=3.0), the cells were harvested via centrifugation and then washed with fresh medium, before being used for the following experiments.

In total, six identical column reactors were employed. Each reactor contained 800 mL of fresh modified f/2 liquid medium supplemented with 10 mM Tris-HCl buffer (pH=8.2) to accurately control the pH during culture. Equal numbers of seed cells were re-inoculated into each of the six-column reactors to an OD_750_ of 1.5. The light intensity was maintained at 80±5 μmol m^-2^ s^-1^. The six algal cultures were first aerated with air enriched with 5% CO_2_ (“High CO_2_” conditions, or HC) for one hour. After the pre-adaption phase, three of the algal cultures proceeded under HC as the control condition, whereas the other three were switched to aeration with 0.01% CO_2_ (“Low CO_2_” conditions, or LC) (Brueggeman et al., 2012; Fang et al., 2012) for CCM induction (**Fig. S1**). After being transferred to the designated conditions, cell aliquots were taken at 0 and 24 h from each column via a syringe for the profiling of DNA methylation, histone modification, chromatin accessibility and chromatin interaction.

For the mutants, the cells were cultured in liquid cultures of the f/2 medium and maintained under light at 50 μmol photos m^-2^ s^-1^ at 25 °C blowing air or under 5% CO_2_. The growth curve was generated via cell counting, with three replicates for each strain. After 6 days, the cells were harvested via centrifugation and stored at -80 °C, before being used for the following experiments.

### RNA-Seq profiling and data analysis

The RNA-Seq datasets of LC experiments were retrieved from the NCBI GEO database with accession number GSE55861. For the mutants, a total of 2 μg of RNA per sample was used as input material for the RNA sample preparations. Sequencing libraries were constructed as previously described (Wei et al., 2019). Pair-end sequencing (2*150 bp) of each sample was performed on the Illumina HiSeq platform (Illumina, USA).

The RNA-seq datasets were analyzed mainly via the nf-core/rnaseq pipeline v1.4.2 (https://github.com/nf-core/rnaseq). In brief, the raw reads were quality-controlled via Trim Galore (https://github.com/FelixKrueger/TrimGalore). The high-quality read pairs were aligned to the *N. oceanica* IMET1 genome (Gong et al., 2020) with STAR (Dobin et al., 2013) with modified parameters to limit intron lengths (“--alignIntronMin 20 --alignIntronMax 3000”) and duplicates marked via Picard (http://broadinstitute.github.io/picard/). Then, the abundance of expression was quantified via featureCounts (Liao et al., 2013) and StringTie (Pertea et al., 2016). Then, scripts from Trinity (Grabherr et al., 2011) were used to generate a gene expression matrix via the RSEM2 method and with TMM normalization to account for library size variation between samples. After that, TPM (transcripts per kilobase million) values were averaged among replicates. For LC experiments, raw counts of HC (0 h) and LC (VLC_24h) samples from featureCounts (Liao et al., 2013) were used for differential gene expression analysis with edgeR (Robinson et al., 2010) with an FDR ≤ 0.001 and a minimum fold change > 2.

### ChIP-Seq profiling and data analysis

*N. oceanica* cells (1-2 million cells/mL) from 800 mL cultures were fixed under vacuum for 30 min with 1% formaldehyde (Sigma, USA) at room temperature. After fixation, the cells were incubated at room temperature for 10 min under vacuum with 0.15 M glycine. Nuclei were isolated as described previously (Wei and Xu, 2018), and nuclei from 1 g of fixed material were used for each round of ChIP. The isolated nuclei were resuspended in 1 mL of sonication buffer (10 mM potassium phosphate, pH 7.0, 0.1 mM NaCl, 0.5% sarkosyl, and 10 mM EDTA), and the chromatin was sheared by sonication with Covaris S220 to achieve an average fragment size of around 300-1000 bp. The sonicated sample was mixed with 100 µL of 10% Triton X-100, and 50 µL of this mixture was saved as the input sample. The rest of the sheared chromatin was mixed with an equal volume of IP buffer (50 mM HEPES, pH 7.5, 150 mM NaCl, 5 mM MgCl2, 10 µM ZnSO4, 1% Triton X-100, 0.05% SDS) and incubated with anti-H3K9Ac (Abcam, UK; ab10812), anti-H3K27Ac (Millipore, Germany; 07-360), anti-H3K4me2 (Abcam, UK; ab7766) or anti-crotonyllysine (PTM BioLab, USA; PTM-501) antibodies. After overnight incubation at 4 °C, 50 µL of protein A/G magnetic beads (Millipore, Germany) were added and incubated at 4 °C for 2 h. The beads were washed at 4 °C as follows: 3× with IP buffer, 1× with IP buffer containing 500 mM NaCl, and 1× with LiCl buffer (0.25 M LiCl, 1% NP-40, 1% deoxycholate, 1 mM EDTA, 10 mM Tris pH 8.0) for 5 min each. Chromatin retained on beads was incubated in 200 µL of elution buffer (50 mM Tris, pH 8.0, 200 mM NaCl, 1% SDS, and 10 mM EDTA) at 65 °C for 30 min, followed by proteinase K treatment at 65 °C for 6 h. DNA was extracted with a Qiagen kit (Qiagen, Germany), and all subsequent end-repair, A-tailing, adaptor ligation, and library amplification steps were performed following a standard protocol (Illumina, USA). The final library was sequenced on an Illumina HiSeq 2000 instrument with 2×150-bp reads.

Additionally, we used three antibodies for histone modifications (H3K9me3, H3K27me3, and H2A.Z) to profile the genome-wide distribution in *N. oceanica*. Among them, H3K9me3 and H3K27me3 are present in *P. tricornutum* (Veluchamy et al., 2015) and H2A.Z was detected in *Arabidopsis* (Gómez-Zambrano et al., 2019). However, they were not effectively enriched via ChIP in *N. oceanica* (ChIP-Seq data were deposited under the Sequence Read Archive with accessions SRR16288281-SRR16288288), which suggests that the histone codes of *N. oceanica* are different from those of diatoms and *Arabidopsis*.

The ChIP-Seq datasets were analyzed mainly via the nf-core/chipseq pipeline v1.2.1 (https://github.com/nf-core/chipseq), with minor modifications. In brief, the reads were trimmed via Trim Galore (https://github.com/FelixKrueger/TrimGalore), then aligned against the reference genome via BWA (Li and Durbin, 2009). The mapped reads were duplication marked via Picard (http://broadinstitute.github.io/picard/) and quality controlled via phantompeakqualtools (Landt et al., 2012) and deepTools (Ramírez et al., 2016). Peak calling was performed via MACS2 (Zhang et al., 2008) with the “--narrow_peak” flag and input samples as controls, followed by peak annotation (relative to gene features) via annotatePeaks.pl from HOMER (Heinz et al., 2010). For each histone modification, consensus peaks were consolidated via BEDTools (Quinlan and Hall, 2010) and then quantified via featureCounts (Liao et al., 2013), followed by differential binding analysis via DESeq2 (Love et al., 2014) (FDR < 0.01). The promoter of each gene is defined as the fragment from 1 kbp upstream of this gene to the TSS.

### MNase-Seq library construction and sequencing data analysis

*Nannochloropsis* nuclei were isolated and digested according to Dai’s method (Dai et al., 2017). Briefly, *N. oceanica* cells (1-2 million cells/mL) from 800 mL cultures were fixed under vacuum for 30 min with 1% formaldehyde (Sigma, USA) at room temperature. After fixation, the cells were incubated at room temperature for 10 min under vacuum with 0.15 M glycine (Wei and Xu, 2018). The cells were collected via centrifugation at 2500×g and ground under liquid nitrogen in a mortar and pestle. Subsequently, the ground cell powder was resuspended in hypertonic buffer A (50 mM HEPES [pH 7.5], 1 mM EDTA [pH 8.0], 150 mM NaCl, 1% Triton X-100, and 1x protease inhibitor cocktail (Roche, Switzerland)), and shaken for 30 min at 4 °C. Solutions were filtered through a Miracloth (Calbiochem, Germany) into fresh 50 mL tubes. Nuclei were collected by centrifugation at 4 °C for 20 min at 4000×g. The pellets were washed twice in buffer A and then resuspended in buffer D (10% sucrose, 50 mM Tris-HCl [pH 7.5], 25 mM MgCl_2_, and 1 mM CaCl_2_). Chromatin was fragmented by incubation for 10 min at 37 °C with 2 units of micrococcal nuclease (Sigma, USA). The reactions were stopped with the addition of 5 mM EDTA. Mononucleosome-sized fragments were gel-purified (Petesch and Lis, 2008). The nucleosomal population was subsequently subjected to paired-end high-throughput sequencing.

The MNase-seq datasets were analyzed mainly via the nf-core/mnaseseq pipeline v1.0 (https://github.com/nf-core/mnaseseq). In brief, the raw reads of the MNase-seq data were quality controlled via Trim Galore (https://github.com/FelixKrueger/TrimGalore), after which high-quality clean reads were aligned to the reference genome (IMET1v2) via Bowtie2 (Langmead and Salzberg, 2012). Unpaired and discordant alignments were discarded, and duplicates were marked via Picard (http://broadinstitute.github.io/picard/). The alignments from the replicates were merged and then quality controlled via deepTools (Ramírez et al., 2016). The nucleosome positions, occupancy levels, and differential occupancies between the LC and HC samples were calculated via DANPOS2 (Chen et al., 2013).

### Chromatin state analysis

ChromHMM (Ernst and Kellis, 2017) was used to integrate MNase-seq data and ChIP-seq data of H3k9ac, H3k27ac, Kcr, and H3k4me2 to discover the major re-occurring combinatorial and spatial patterns of marks. Several models were trained (“LearnModel” mode) in parallel mode with the number of states ranging from 3 states to 9 states and via the “CompareModels” mode of ChromHMM (Ernst and Kellis, 2017). The model of five chromatin states was selected, and then the chromatin state of each gene was defined as the chromatin state at the TSS. From HC to LC, the transition of the chromatin states of genes was summarized via custom scripts. GO and pathway enrichment analyses of the gene sets of different transition types were performed via services provided by the NanDeSyn database (Gong et al., 2020).

### Generation of targeted knockout *N. oceanica* mutants via CRISPR

A modular CRISPR/Cas9 toolbox system (pNOC-ARS-CRISPR; (Poliner et al., 2018)) was used to construct the CRISPR plasmids. Briefly, gRNAs were designed via the CHOPCHOP platform (http://chopchop.cbu.uib.no). The CRISPR/Cas9 episome was constructed as described by Poliner et al. (2018), and the episome system was transformed into *N. oceanica* via an electrophoresis protocol (Wang et al., 2016). The target sites of the mutants and WT were amplified via genome PCR and the PCR products were detected via Sanger sequencing.

#### Photosynthesis parameters monitoring

The chlorophyll fluorescence of the mutants and WT (as a control) were measured via a pulse-amplitude modulated fluorometer (Image PAM, Walz, Effeltrich, Germany) after 20 min of dark treatment. The PSII maximum quantum yield (Fv/Fm) was measured according to previous reports (Maxwell and Johnson, 2000; Wei et al., 2017).

#### ChIP-Seq sequencing

For the cross-linking reaction, the mutants and WT (as a control) were incubated with 37% formaldehyde as previously described (Wei and Xu, 2018), and the cells were incubated with an anti-H3K4me2 antibody (Cell Signaling Technology, USA; #9725). The ChIP-Seq library was constructed by Novogene Corporation (Beijing, China). Library quality was assessed on an Agilent Bioanalyzer 2100 system. Subsequently, pair-end sequencing (2*150) of each sample was performed on an Illumina platform (Illumina, USA).

### Overexpression of NoPMA2

To construct a vector for NoPMA2 overexpression, the NoPMA2 sequence from *N. oceanica* IMET1 was amplified from cDNA via PCR (Primer F: 5’-acctctttttaaattggtaccatggttgttgtcgacgagtcc-3’; Primer R: 5’-tctcttccggtaggggtacctcagaccataatcgagatatcagat-3’). The PCR-amplified sequence was inserted into the *KpnI* digested vector pXJ53-1 (GenBank accession OR338766.1). NoPMA2 was expressed by the endogenous *VCP1* promoter and the *bleR* gene was used as a selection marker. The transformants were cultured in triangular flasks at 50 μmol m^-2^ s^-1^ with shaking at 200 rpm at 25 °C.

### Hi-C sequencing and data analysis

Following the standard protocol described previously with certain modifications (Belton et al., 2012), we constructed Hi-C libraries using *N. oceanica* cells as inputs (under LC and HC conditions). Briefly, microalgal cells were crosslinked with 2% formaldehyde solution at room temperature under vacuum for 30 min. Then, 2.5 M glycine was added to quench the crosslinking reaction for 10 min at room temperature. After being ground with liquid nitrogen and re-suspended with 25 ml of extraction buffer I (0.4 M sucrose, 10 mM Tris-HCl, pH 8, 10 mM MgCl_2_, 5 mM β-mercaptoethanol, 0.1 mM phenylmethylsulfonyl fluoride [PMSF], and 13 units of protease inhibitor), the mixture was further centrifuged at 4000 rpm at 4 °C for 20 min. Re-suspended pellet in extraction buffer II (0.25 M sucrose, 10 mM Tris-HCl, pH 8, 10 mM MgCl2, 1% Triton X-100, 5 mM β-mercaptoethanol, 0.1 mM PMSF, and 13 units of protease inhibitor) was centrifuged at 14,000 rpm and 4 °C for 10 min. The pellet was re-suspended in extraction buffer III (1.7 M sucrose, 10 mM Tris-HCl, pH 8, 0.15% Triton X-100, 2 mM MgCl2, 5 mM β-mercaptoethanol, 0.1 mM PMSF, and 13 units of protease inhibitor) and loaded on the top of an equal amount of clean extraction buffer III, which was then centrifuged at 14,000 rpm for 10 min. The pellet was washed twice in 500 μL ice cold 1x CutSmart buffer and then centrifuged for 5 min at 2,500 g. The nuclei were washed by 0.5 mL of restriction enzyme buffer and solubilized with dilute SDS followed by incubation at 65 °C for 10 min. After quenching the SDS by Triton X-100, overnight digestion was applied to the samples with a 4-cutter restriction enzyme MboI (400 units) at 37 °C on a rocking platform.

The subsequent steps involved marking the DNA ends with biotin-14-dCTP and blunt-end ligation of the cross-linked fragments. The proximal chromatin DNA was re-ligated by a ligation enzyme. The nuclear complexes were reverse crosslinked via incubation with proteinase K at 65 °C. DNA was purified via phenol-chloroform extraction. Biotin was removed from non-ligated fragment ends via T4 DNA polymerase. Ends of sheared fragments by sonication (200-600 base pairs) were repaired by a mixture of T4 DNA polymerase, T4 polynucleotide kinase, and Klenow DNA polymerase. Biotin-labeled Hi-C samples were specifically enriched via streptavidin C1 magnetic beads. After the addition of the A-tails to the fragment ends and subsequent ligation with Illumina paired-end (PE) sequencing adapters, the Hi-C sequencing libraries were amplified via PCR (10-15 cycles) and sequenced on an Illumina HiSeq 2500 platform (2*125 bp).

The Hi-C sequencing datasets were analyzed via the HiC-Pro pipeline v2.9.0 (Servant et al., 2015), which uses Bowtie2 (Langmead and Salzberg, 2012) to map sequencing reads to the reference genome, performs various quality filtering procedures, and normalizes the contact maps via ICE normalization (https://github.com/hiclib/iced). The interaction maps were visualized and compared via HiCPlotter (Akdemir and Chin, 2015). Based on the 10 kb resolution maps, the 3D structure of the genomes was reconstructed via 3DMax v1.0 (Oluwadare et al., 2018), which employs the maximum likelihood algorithm and visualizes via NGLVieweR v1.3.1 (https://github.com/nvelden/NGLVieweR/). The compartments were called via cooltools v0.4.0 (https://github.com/open2c/cooltools). Significant chromatin contacts were identified via FitHiC v1.1.3 (Ay et al., 2014), significant interaction differences were analyzed via HiCcompare v1.8.0 (Stansfield et al., 2018), and chromatin loops were called via Juicer v1.5.6 (Durand et al., 2016). The significant items were retained if supported by both biological replicates.

### WGBS and data analysis

A total of 5.2 μg of genomic DNA was fragmented into 200-300 bp fragments via sonication (Covaris, USA; S220), followed by end-repair and adenylation. The cytosine-methylated barcodes were ligated to sonicated DNA. DNA fragments were treated twice with bisulfite using an EZ DNA Methylation-GoldTM Kit (Zymo Research, USA) before the resulting single-stranded DNA fragments were PCR amplified via KAPA HiFi HotStart Uracil + ReadyMix (2X). Libraries were quantified by a Qubit® 2.0 fluorometer (Life Technologies, USA) and quantitative RT-PCR, and the insert size was assessed on an Agilent Bioanalyzer 2100 system. Clustering of the index-coded samples was performed on a cBot Cluster Generation System via the TruSeq PE Cluster Kit v3-cBot-HS (Illumina, USA) according to the manufacturer’s instructions. After cluster generation, the library preparations were sequenced on an Illumina HiSeq 2500 platform and 125 bp paired-end reads were generated. Image analysis and base calling were performed with the Illumina CASAVA pipeline, and 125 bp paired-end reads were generated.

Bismark software (version 0.16.1) (Krueger and Andrews, 2011) was used to perform alignments of bisulfite-treated reads to the reference genome via default parameters. The results of the methylation extractor were transformed into bigWig format for visualization via the IGV browser (Thorvaldsdóttir et al., 2012). Only cytosines with read support are considered valid, and every valid cytosine is regarded as equal. The bisulfite conversion efficiencies were estimated based on unmethylated λ phage which was spiked in the DNA as a control. Differentially methylated regions (DMRs) were detected via the R/Bioconductor package DMRcaller v1.26.0 (Catoni et al., 2018) with default parameters and by combining multiple methods (“noise_filter”, “neighbourhood” and “bins”). The methylation level of every valid cytosine (MLmc) was calculated by the read count of methylated cytosines (mCs) mapped to their genomic locus divided by the read count of all cytosines mapped to the same locus. The methylation level of a specific genomic fragment (MLgf) was calculated as the average methylation level of all cytosines (and guanines in the reverse strand) of this genomic fragment. The methylation level of a class of genomic components (MLgc) was calculated as the average MLmc of all cytosines belonging to the specific component. This approach is also used to calculate the methylation level of the whole genome (MLwg). To remove experimental artifacts, noise, and any batch effects, MLgc was normalized according to MLwg, which is defined as the normalized methylation level of genomic components (NMLgc) (Xiang et al., 2010).

## Data availability

For ChIP-Seq, MNase-Seq, WGBS, and Hi-C, raw sequencing data for the wild-type *N. oceanica* have been deposited in the Sequence Read Archive under accessions PRJNA769792, PRJNA769749, PRJNA769619 and PRJNA513887 respectively. The RNA-Seq and ChIP-Seq data for the mutants have been deposited as PRJNA769958. These 73 datasets have also been integrated into the NanDeSyn database (Gong et al., 2020) and are accessible for comparative analyses with existing *Nannochloropsis* omics data via a genome browser (http://nandesyn.single-cell.cn/browser).

## Author contributions

Y.G., Q.W., L.S., and J.X. contributed to the conception and design of the research as well as to the writing of the manuscript; L.W. and Q.W. performed the experiments and acquired the data; Q.W., LH.W., N.L., and X.D. created the mutants and profiled the phenotypes; Y.G. performed the bioinformatic analyses; Y.G., Q.W., C.S, Y.X., L.S. and J.X. contributed to the analysis and interpretation of the data. All the authors read and approved the final manuscript.

## Acknowledgment

This work was supported by Synthetic Biology Program from Ministry of Science and Technology of China (2021YFA0909700), National Natural Science Foundation of China (grant numbers 32200448, 32370097, 31741005 and 31900071), Biological Carbon Sequestration Program (ZDRW-ZS-2016-3 and KSZD-EW-Z-017) from Chinese Academy of Sciences, and I201908 from Qingdao Institute of Bioenergy and Bioprocess Technology, Chinese Academy of Sciences.

