## Supplementary material for "Dynamic epigenomic landscape of carbon-concentrating mechanisms in the model industrial oleaginous microalga *Nannochloropsis oceanica*": Fig. S, Table S

**Running title**: Epigenomic regulation of carbon assimilation in microalgae

### Supplemental Tables

**Table S1. Statistics of the *N. oceanica* WGBS data.**

| Sample | Reads (M) | Aligned reads (M) | Total Cs (M) | Methylated CpGs | Methylated CpHs | Methylated CHHs |
|---|---|---|---|---|---|---|
| HC_1 | 3.82 | 2.19 | 70.68 | 19,986 (0.13%) | 18,051 (0.14%) | 49,274 (0.12%) |
| HC_2 | 4.51 | 2.54 | 78.43 | 21,973 (0.13%) | 20,444 (0.15%) | 54,905 (0.12%) |
| HC_3 | 5.71 | 3.05 | 96.21 | 26,746 (0.13%) | 24,763 (0.15%) | 65,806 (0.12%) |
| LC_1 | 7.72 | 4.91 | 157.43 | 50,436 (0.15%) | 46,008 (0.16%) | 121,692 (0.13%) |
| LC_2 | 6.92 | 3.88 | 121.28 | 33,944 (0.13%) | 31,684 (0.15%) | 85,691 (0.12%) |
| LC_3 | 6.03 | 3.22 | 101.14 | 28,492 (0.13%) | 26,069 (0.15%) | 71,031 (0.12%) |

**Table S2. Read statistics and quality control of *N. oceanica* ChIP-Seq data.** Valid read pairs are high-quality mapped read pairs after filtering out duplicates, non-primary alignments, multiple alignments, mapped reads with > 4 mismatches, read pairs > 2 kb insert size, and read pairs not in forward-reverse orientation. NSC: normalized strand cross-correlation coefficient; RSC: relative strand cross-correlation coefficient.

| Sample | Raw read pairs (M) | Mapped read pairs (M) | Valid read pairs (M) | NSC | RSC |
|---|---|---|---|---|---|
| HC_H3K27ac_R1 | 5.49 | 5.47 | 3.76 | 1.14 | 1.09 |
| HC_H3K27ac_R2 | 8.09 | 8.07 | 5.94 | 1.13 | 1.12 |
| HC_H3K4me2_R1 | 16.06 | 16.06 | 8.61 | 1.04 | 1.34 |
| HC_H3K4me2_R2 | 11.91 | 11.90 | 7.39 | 1.04 | 1.33 |
| HC_H3K9ac_R1 | 10.06 | 8.33 | 5.53 | 1.10 | 1.18 |
| HC_H3K9ac_R2 | 17.86 | 15.23 | 8.74 | 1.04 | 1.24 |
| HC_Kcr_R1 | 6.01 | 6.00 | 5.03 | 1.20 | 1.14 |
| HC_Kcr_R2 | 12.95 | 12.95 | 10.81 | 1.19 | 1.15 |
| LC_H3K27ac_R1 | 4.51 | 4.50 | 2.78 | 1.12 | 1.11 |
| LC_H3K27ac_R2 | 6.68 | 6.68 | 4.73 | 1.11 | 1.15 |
| LC_H3K4me2_R1 | 22.80 | 22.79 | 11.71 | 1.04 | 1.35 |
| LC_H3K4me2_R2 | 15.46 | 15.45 | 9.44 | 1.04 | 1.46 |
| LC_H3K9ac_R1 | 14.27 | 13.40 | 10.06 | 1.22 | 1.16 |
| LC_H3K9ac_R2 | 25.17 | 23.03 | 18.01 | 1.20 | 1.10 |
| LC_Kcr_R1 | 3.64 | 3.64 | 2.96 | 1.14 | 1.17 |
| LC_Kcr_R2 | 4.38 | 4.38 | 3.59 | 1.15 | 1.22 |
| Input_HC_R1 | 10.27 | 8.70 | 5.32 | - | - |
| Input_HC_R2 | 20.82 | 13.81 | 8.34 | - | - |
| Input_LC_R1 | 11.49 | 10.05 | 6.39 | - | - |
| Input_LC_R2 | 11.28 | 10.13 | 6.59 | - | - |

**Table S3. Statistics of peak calling and annotation for *N. oceanica* ChIP-Seq data.** FRiP: fraction of reads in peaks.

| Sample | # peaks | FRiP | # peaks in promoter-TSS | # peaks in intergenic | # unique genes |
|---|---|---|---|---|---|
| HC_H3K27ac_R1 | 6,007 | 0.68 | 4,529 | 46 | 5,472 |
| HC_H3K27ac_R2 | 6,006 | 0.70 | 4,486 | 44 | 5,457 |
| HC_H3K4me2_R1 | 9,957 | 0.54 | 6,170 | 71 | 7,486 |
| HC_H3K4me2_R2 | 10,261 | 0.56 | 6,361 | 67 | 7,664 |
| HC_H3K9ac_R1 | 5,941 | 0.54 | 4,486 | 27 | 5,454 |
| HC_H3K9ac_R2 | 5,767 | 0.38 | 4,279 | 21 | 5,283 |
| HC_Kcr_R1 | 6,124 | 0.79 | 4,552 | 48 | 5,563 |
| HC_Kcr_R2 | 6,159 | 0.80 | 4,570 | 45 | 5,569 |
| LC_H3K27ac_R1 | 5,640 | 0.63 | 4,255 | 25 | 5,173 |
| LC_H3K27ac_R2 | 5,587 | 0.64 | 4,149 | 24 | 5,129 |
| LC_H3K4me2_R1 | 10,387 | 0.62 | 6,309 | 77 | 7,768 |
| LC_H3K4me2_R2 | 9,816 | 0.61 | 5,949 | 59 | 7,465 |
| LC_H3K9ac_R1 | 6,180 | 0.74 | 4,713 | 29 | 5,643 |
| LC_H3K9ac_R2 | 6,176 | 0.73 | 4,703 | 28 | 5,649 |
| LC_Kcr_R1 | 5,795 | 0.67 | 4,279 | 22 | 5,300 |
| LC_Kcr_R2 | 5,772 | 0.69 | 4,255 | 22 | 5,283 |

**Table S4. Associations between gene transcription and histone marks in *N. oceanica*.** $P_+$: the number of up-regulated peaks; $P_-$: the number of down-regulated peaks; $G_+$: the number of up-regulated genes; $G_-$: the number of down-regulated genes.

| | **$P_+$** | **$(P_+G_+)/(P_+G_-)$** | **$P_-$** | **$(P_-G_-)/(P_-G_+)$** |
|---|---|---|---|---|
| H3K9ac | 774 | 162/54=3.00 | 1,049 | 239/168=1.42 |
| H3K27ac | 958 | 299/56=5.34 | 1,211 | 461/123=3.75 |
| Kcr | 1,163 | 312/79=3.95 | 1,289 | 454/141=3.22 |
| H3K4me2 | 145 | 12/68=0.18 | 118 | 9/58=0.16 |

**Table S5. Statistics of *N. oceanica* MNase-Seq data.** Reads were counted in million read pairs.

| Sample | Raw reads | Clean reads | Mapped reads | Duplication (%) | Nucleosomes | Nucleosomes (merged) |
|---|---|---|---|---|---|---|
| HC_1 | 14.28 | 14.00 | 13.64 | 18.97 | 81,200 | 81,031 |
| HC_2 | 8.25 | 8.13 | 7.94 | 15.18 | 81,719 | |
| LC_1 | 6.85 | 6.76 | 6.66 | 17.03 | 87,358 | 87,845 |
| LC_2 | 11.83 | 11.71 | 11.50 | 19.60 | 87,722 | |

**Table S6. Differential expression of the *N. oceanica* DEGs annotated with GO:0005840 and harboring DPNs.**

| Gene | Description | $log_2FC$ |
|---|---|---:|
| NO01G04980 | Ribosomal protein s27a | -2.13 |
| NO02G02810 | 60s ribosomal protein l24 | -1.55 |
| NO02G04340 | 60s ribosomal protein l37a | -2.49 |
| NO03G00290 | 60S ribosomal protein L36 | -1.92 |
| NO03G01890 | Ribosomal protein L35 | -1.60 |
| NO03G04360 | 60s ribosomal protein l11 | -1.86 |
| NO03G05070 | 60s acidic ribosomal protein | -2.61 |
| NO03G05230 | Ribosomal protein L38 | -2.25 |
| NO04G01930 | 60s acidic ribosomal protein p0 | -2.01 |
| NO04G01970 | Ubiquitin | -1.94 |
| NO05G00730 | Ribosomal protein S7, conserved site | -1.67 |
| NO06G03940 | 40s ribosomal protein s3-3 | -2.14 |
| NO08G00250 | 60s ribosomal protein l31 | -2.21 |
| NO09G01130 | Ribosomal protein L18e/L15P | -1.56 |
| NO09G01420 | 40S ribosomal protein S26 | -2.25 |
| NO09G02240 | Ribosomal protein L18e/L15P | -1.98 |
| NO09G03120 | Ribosomal protein L15 | -2.14 |
| NO10G00250 | Ribosomal protein L2 family | -1.63 |
| NO15G01560 | Translation protein SH3 | -1.83 |
| NO17G00120 | 40S ribosomal protein S27 | -2.74 |
| NO17G00590 | 40S ribosomal protein SA | -2.15 |
| NO18G00920 | Small subunit ribosomal protein S7e | -2.11 |
| NO19G00670 | ribosomal protein S13A | -1.49 |
| NO20G00720 | 60s ribosomal protein l13 | -1.68 |
| NO21G01100 | Ribosomal protein l27 | -1.99 |
| NO22G01100 | Small subunit ribosomal protein S11e | -1.64 |
| NO28G00080 | ribosomal protein L23AB | -2.69 |
| NO28G00190 | ubiquitin 8 | -1.92 |
| NO28G00930 | Ribosomal protein L7A/L8 | -1.69 |

**Table S7. Differential expression of the *N. oceanica* genes annotated with GO:0005840 and carrying chromatin state transitions of CS5->CS4.**

| Gene | Description | $log_2FC$ |
|---|---|---|
| NO01G04980 | Ribosomal protein s27a | -2.13 |
| NO02G02810 | 60s ribosomal protein l24 | -1.55 |
| NO02G03530 | 60s ribosomal protein l13a | -2.16 |
| NO02G04340 | 60s ribosomal protein l37a | -2.49 |
| NO02G04560 | Ribosomal protein S19e family protein | -2.05 |
| NO03G01740 | 40S ribosomal protein S21 | -2.99 |
| NO03G04360 | 60s ribosomal protein l11 | -1.86 |
| NO04G01970 | Ubiquitin | -1.94 |
| NO05G00730 | Ribosomal protein S7, conserved site | -1.67 |
| NO05G03520 | 40s ribosomal protein s15a | -2.19 |
| NO06G01640 | 60s ribosomal protein l9 | -2.39 |
| NO06G02130 | Ubiquitin-40S ribosomal protein S27a | -1.98 |
| NO08G00250 | 60s ribosomal protein l31 | -2.21 |
| NO10G02500 | 60s ribosomal protein l44 | -1.96 |
| NO10G03120 | 40S ribosomal protein S3a | -1.95 |
| NO15G01560 | Translation protein SH3 | -1.83 |
| NO17G00120 | 40S ribosomal protein S27 | -2.74 |
| NO17G00590 | 40S ribosomal protein SA | -2.15 |
| NO18G00920 | Small subunit ribosomal protein S7e | -2.11 |
| NO19G00670 | ribosomal protein S13A | -1.49 |
| NO20G02070 | Ribosomal protein l32 | -1.85 |
| NO21G01530 | 40s ribosomal protein s15 | -2.42 |
| NO25G01590 | 60S ribosomal protein L3 | -2.46 |
| NO28G00080 | ribosomal protein L23AB | -2.69 |

**Table S8. Statistics of *N. oceanica* Hi-C sequencing data.**

| Sample | Sequenced read pairs | Mapped read pairs | Valid interaction pairs |
|---|---|---|---|
| HC_1 | 319,638,650 | 218,602,866 | 210,057,902 |
| HC_2 | 285,983,030 | 193,194,555 | 181,268,189 |
| LC_1 | 300,631,182 | 209,727,741 | 203,830,548 |
| LC_2 | 278,116,295 | 178,260,924 | 165,418,123 |

**Table S9. Epigenetic patterns of key components of the CCM.**

| Symbol | Gene | H3K9ac | H3K27ac | Kcr | H3K4me2 | DPNs | $log_2FC$ |
|---|---|---|---|---|---|---|---|
| NoCA1 | NO04G04510 | * | * | * | * | * | -0.54 |
| NoCA2 | NO01G01240 | Up | * |  | * | * | 1.19 |
| NoCA3 | NO23G01760 | * | * |  | * | * | -0.45 |
| NoCA4 | NO10G03400 | * | * | * | * | * | 0.34 |
| NoCA5 | NO20G00630 | Up | Up | Up | * | * | 2.59 |
| NoBCT1 | NO08G01230 | Up | Up | * | * (Far) | * | 1.27 |
| NoBCT2 | NO01G02900 | * | * | * | * | Y | -0.71 |
| NoPEPC | NO11G00760 | Down | Down | * | * | * | -1.64 |
| NoPEPCK | NO07G03330 | Up | * | * | * | * | 5.81 |
| NoPPDK1 | NO26G01010 | * | Down | * | * | * | 3.20 |
| NoPPDK2 | NO12G00860 | * | * | * | Down (Far) | * | 3.14 |
| NoME2 | NO26G00480 | * | Down | Down | * | * | -2.11 |
| NoME1 | NO06G00510 | Up | * | * | * (Near) | * | 0.83 |
| NoMDH | NO09G03170 | * | * | * | * | * | -0.42 |
| NoGS | NO02G03600 | * | * | * | * | * | -0.62 |
| NoOAT | NO16G01090 | * | * | * | * | * | 0.00 |
| NoCPS | NO01G04780 | * | * | * | * (Close) | * | -2.15 |
| NoOTC | NO04G04600 | * | * | * | * | * | -0.95 |
| NoASS | NO05G00910 | * | * | * | * (Near) | Y | -1.09 |
| NoASL | NO08G03940 | * | * | * | * (Close) | * | -0.25 |
| NoARG | NO16G01720 | * | * | * | * | * | 0.33 |
| NoODC | NO05G00760 | Down | * | * | * (Near) | Y | -2.44 |

### Supplemental Figures

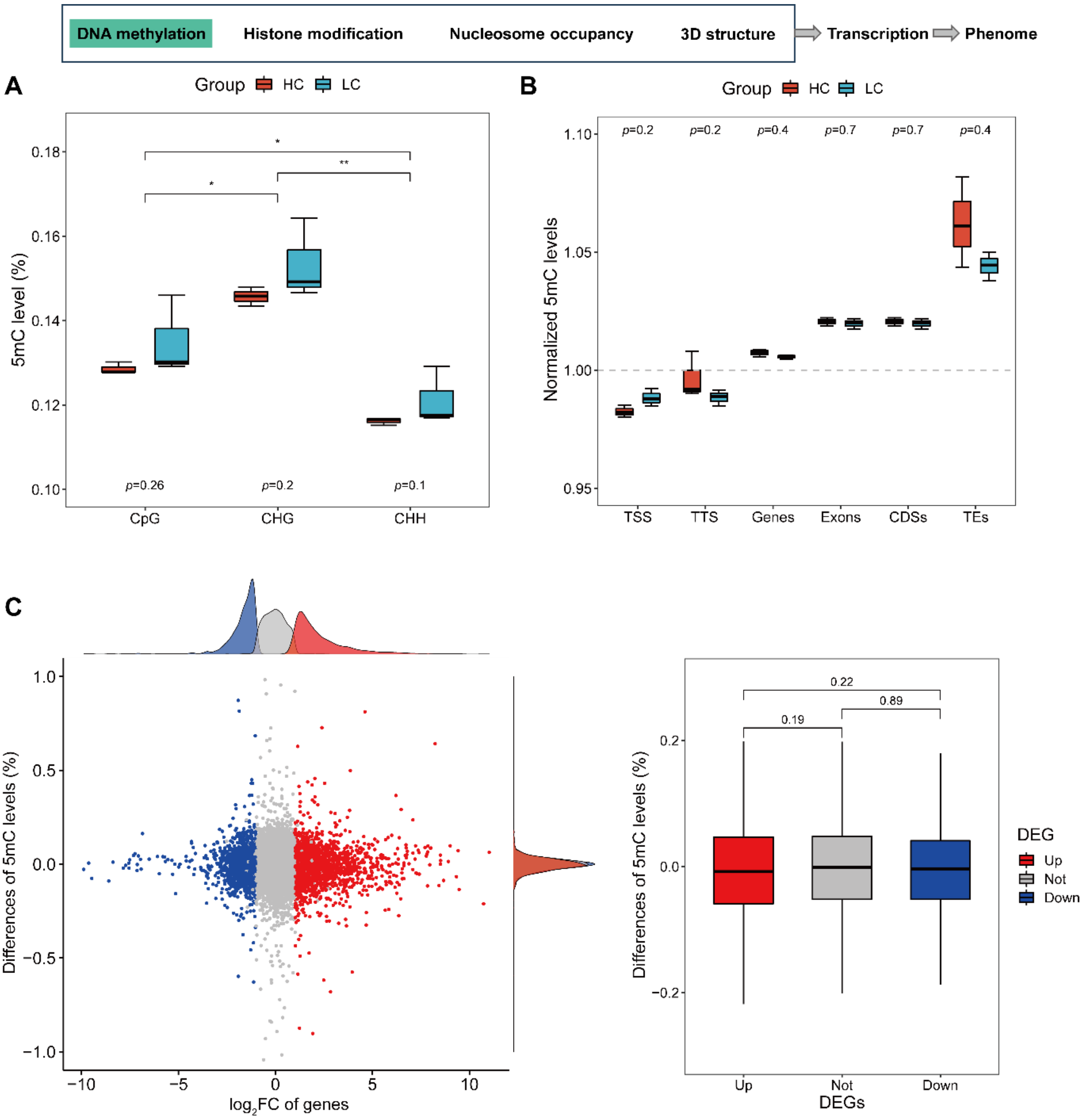


**Figure S1. Characteristics of DNA 5mC methylation in *N. oceanica* under LC stress. (A)** Comparison of 5mC levels in different contexts. (**B**) Normalized 5mC levels in different genomic components. TSS: -1500 bp to TSS; TTS: TTS to +1500 bp.

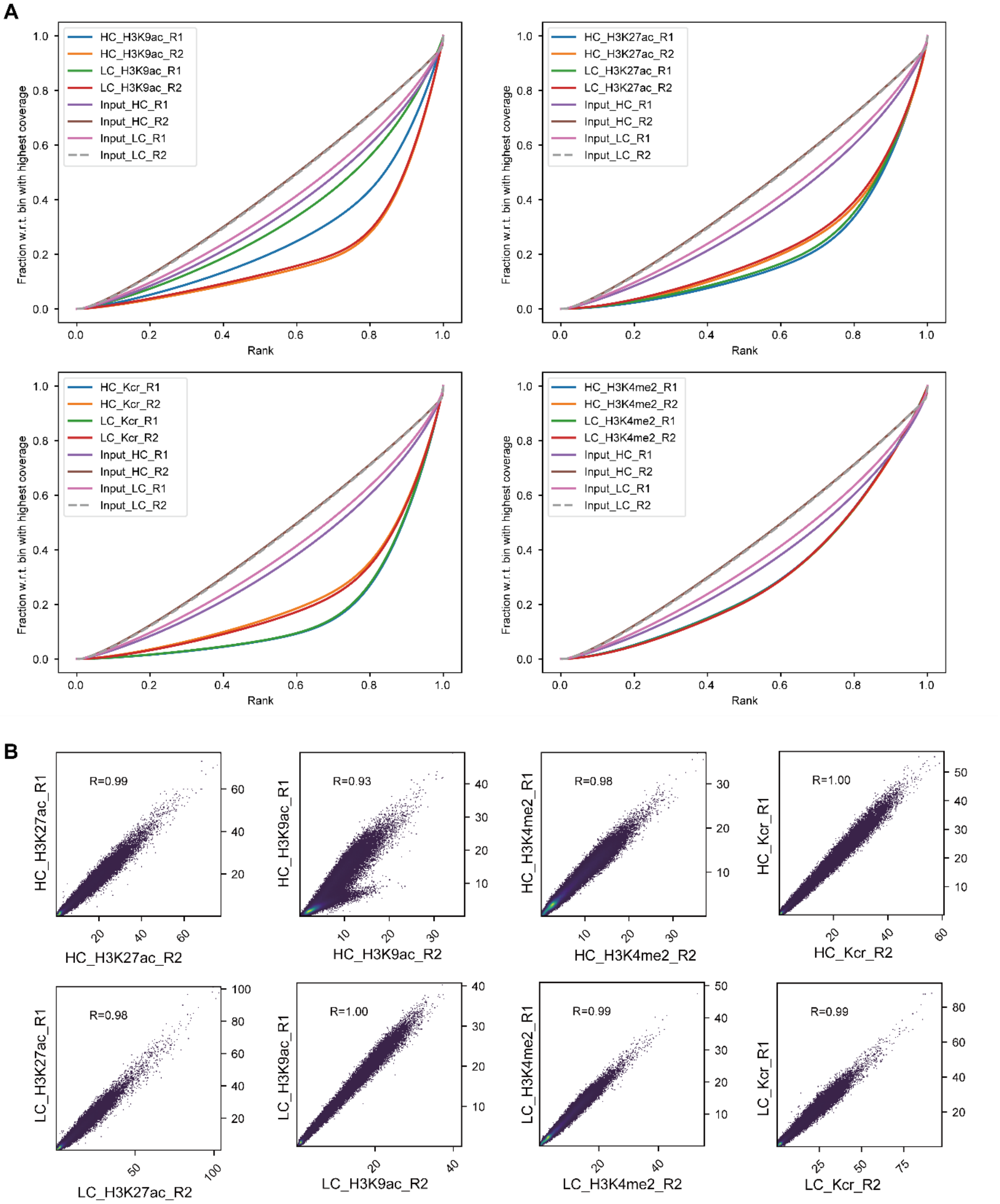


**Figure S2. Quality control for the *N. oceanica* ChIP-Seq datasets.** (**A**) Fingerprint plots showing the enrichment of antibody-treated samples during the ChIP-Seq experiments. (**B**) Sample correlation between biological replicates for ChIP-Seq data. Each point represents a 200 bp non-overlapping bin of the genome; the *x*- and *y*-axes denote the number of fragments in the corresponding samples; the Pearson correlation coefficient between biological replicates is calculated and shown in each panel.

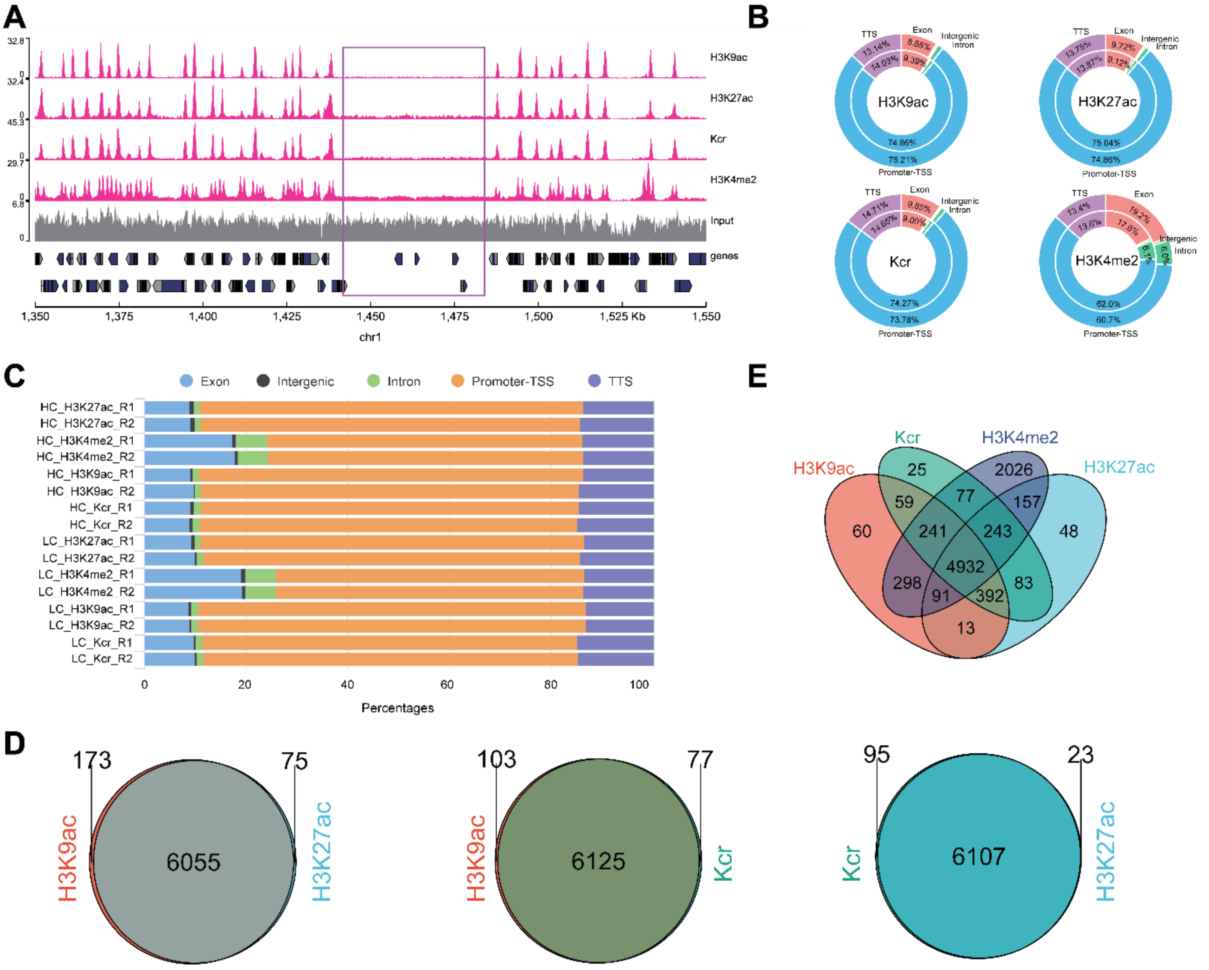


**Figure S3. Overlap between ChIP-Seq peaks and genomic features and between different histone modifications in *N. oceanica*.** (**A**) Patterns of histone marks in gene-poor and gene-rich regions among parts of Chr1. The Y-axis represents the number of reads in a 100 bp interval. (**B**) Proportion of peaks assigned to different genomic features. Most of the peaks reside in the promoter-TSS regions. The "Promoter-TSS" region is defined as the region comprising -1 kbp to +100 bp, and the "TTS" region is defined as the region ranging from -100 bp to +1 kbp. (**C**) Proportion of peaks assigned to different genomic features. Most of the peaks reside in the promoter-TSS regions. (**D**) Venn diagram showing the overlap of detected histone peaks for H3K9ac, H3K27ac, and Kcr. (**E**) Venn diagram showing the overlap of histone-marked genes for H3K4me2, H3K9ac, Kcr and H3K27ac.

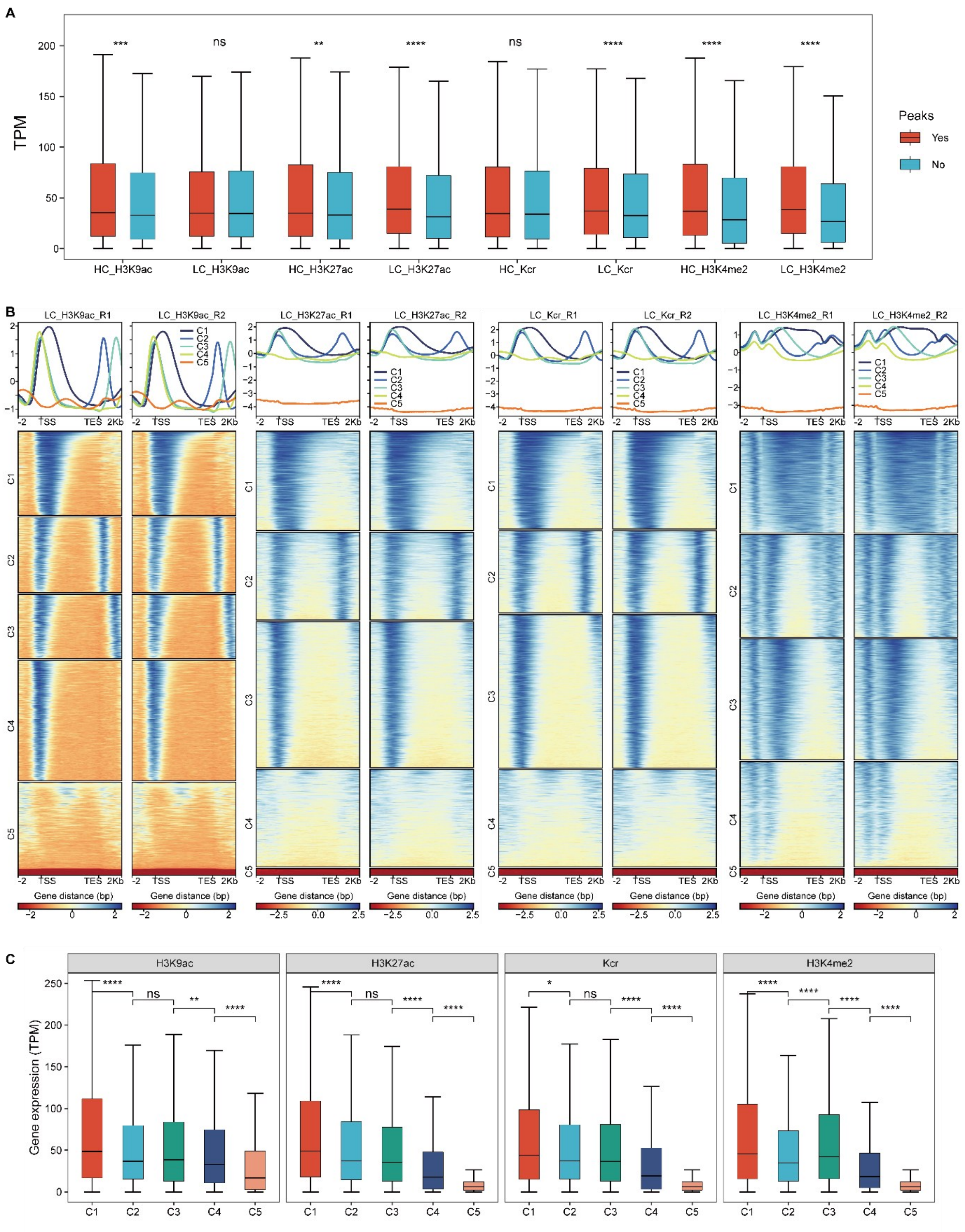


**Figure S4. Links between histone marks and gene expression in *N. oceanica*. (A)** Comparison of gene expression between histone-marked genes and non-marked genes. (**B**) Clustering of ChIP-Seq signals around genes for LC conditions. (**C**) Expression comparison of genes in different clusters.

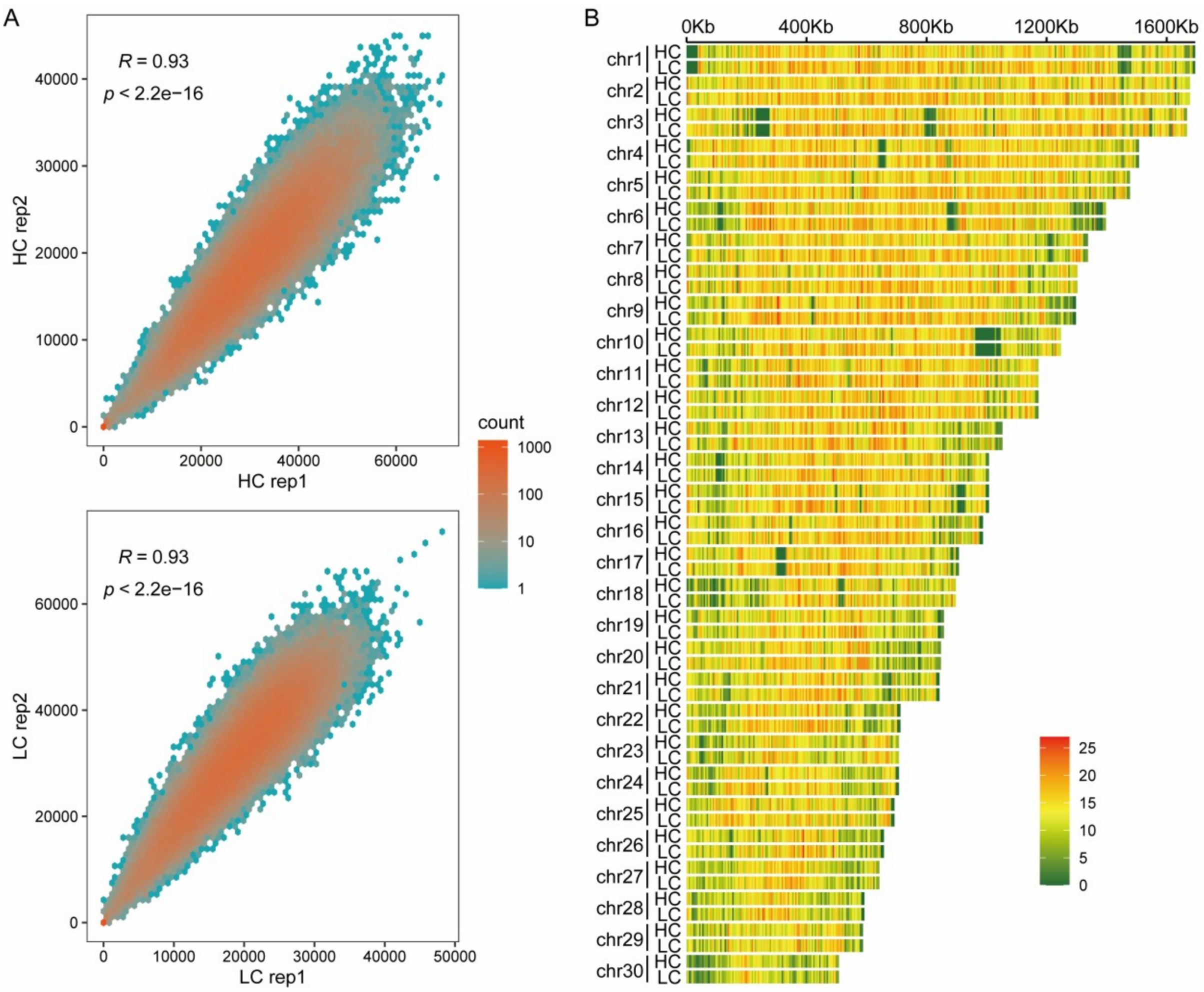


**Figure S5. Identification of nucleosome positioning with MNase-seq for HC and LC samples in *N. oceanica*.** (**A**) Pearson correlations of raw read counts within a bin size of 500 bp between two biological replicates in HC and LC samples, showing a high coefficient value of 0.93. (**B**) Nucleosome density distribution for HC and LC samples within a range of 5 kb on each chromosome. The replicates were combined.

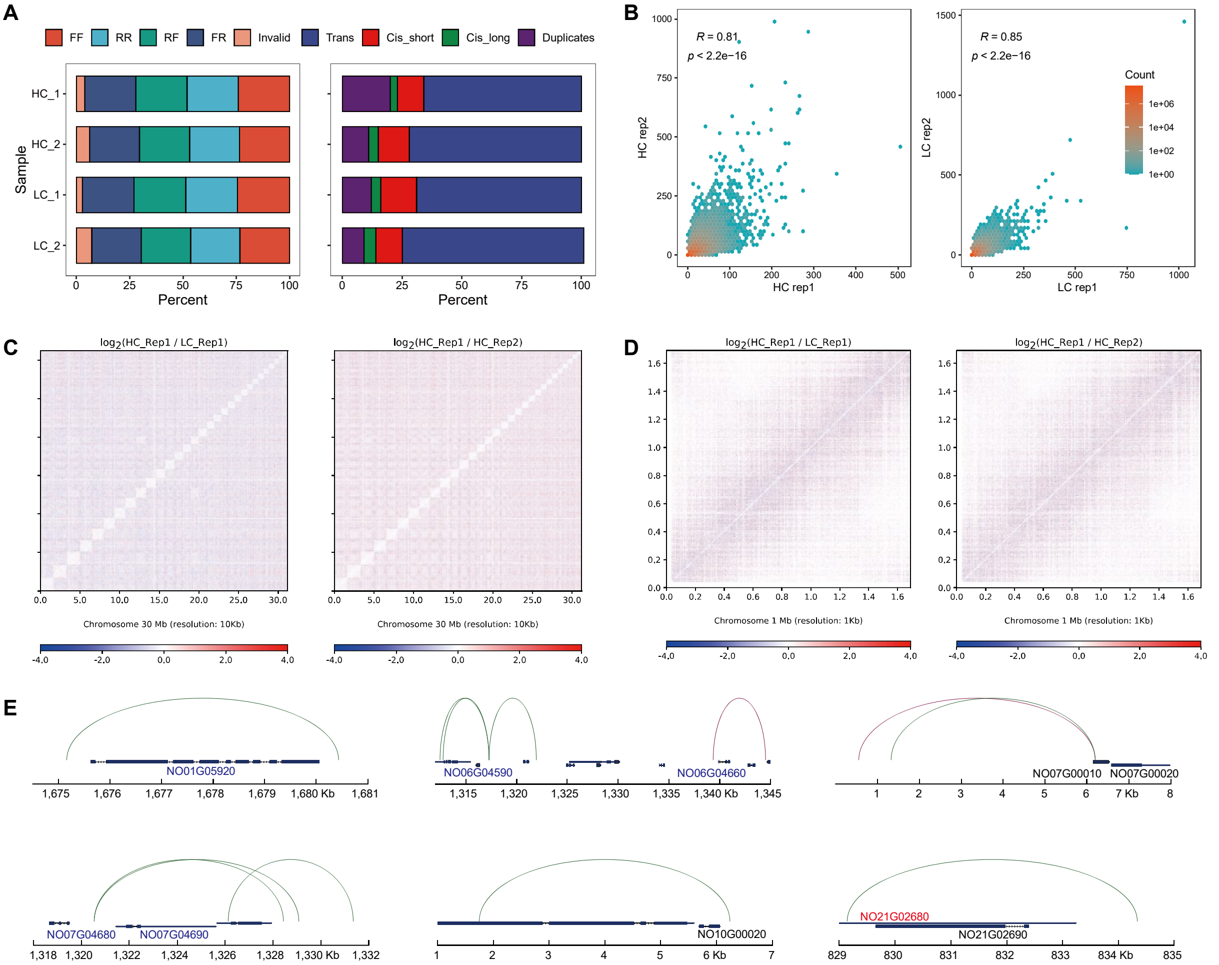


**Figure S6. Quality control and sample comparison for the Hi-C datasets in *N. oceanica*.** (**A**) Quality control of the Hi-C sequencing data. FF: valid interaction read pairs in the forward-forward strand; RR: valid interaction read pairs in the reverse-reverse strand; RF: valid interaction read pairs in the reverse-forward strand; FR: valid interaction read pairs in the forward-reverse strand; Invalid: invalid interaction pairs, including dumped pairs, self-cycle pairs, relegation pairs, single-end pairs and dangling pairs. FF+RR+RF+FR = valid pairs. Trans: trans-contacts; Cis_short: cis-contacts in short-range (< 20 kbp); Cis_long: cis-contacts in long-range (> 20 kbp); Duplicates: duplicated read pairs. (**B**) Hexagonal heatmap plot showing the correlation of chromatin interaction frequencies among biological replicates. The Pearson correlation coefficient was used. The interaction matrixes were in 1 kb resolution. (**C&D**) Hi-C difference matrix between the HC and LC conditions and the two replicates for the whole genome (**C**) and chromosome 1 (**D**). (**E**) Significant chromatin contacts overlapping with gene promoters were visualized. Brown arcs denote significant chromatin contacts in HC samples, whereas green arcs represent those in LC samples. The upregulated DEGs are highlighted in red, whereas the downregulated DEGs are shown in blue.

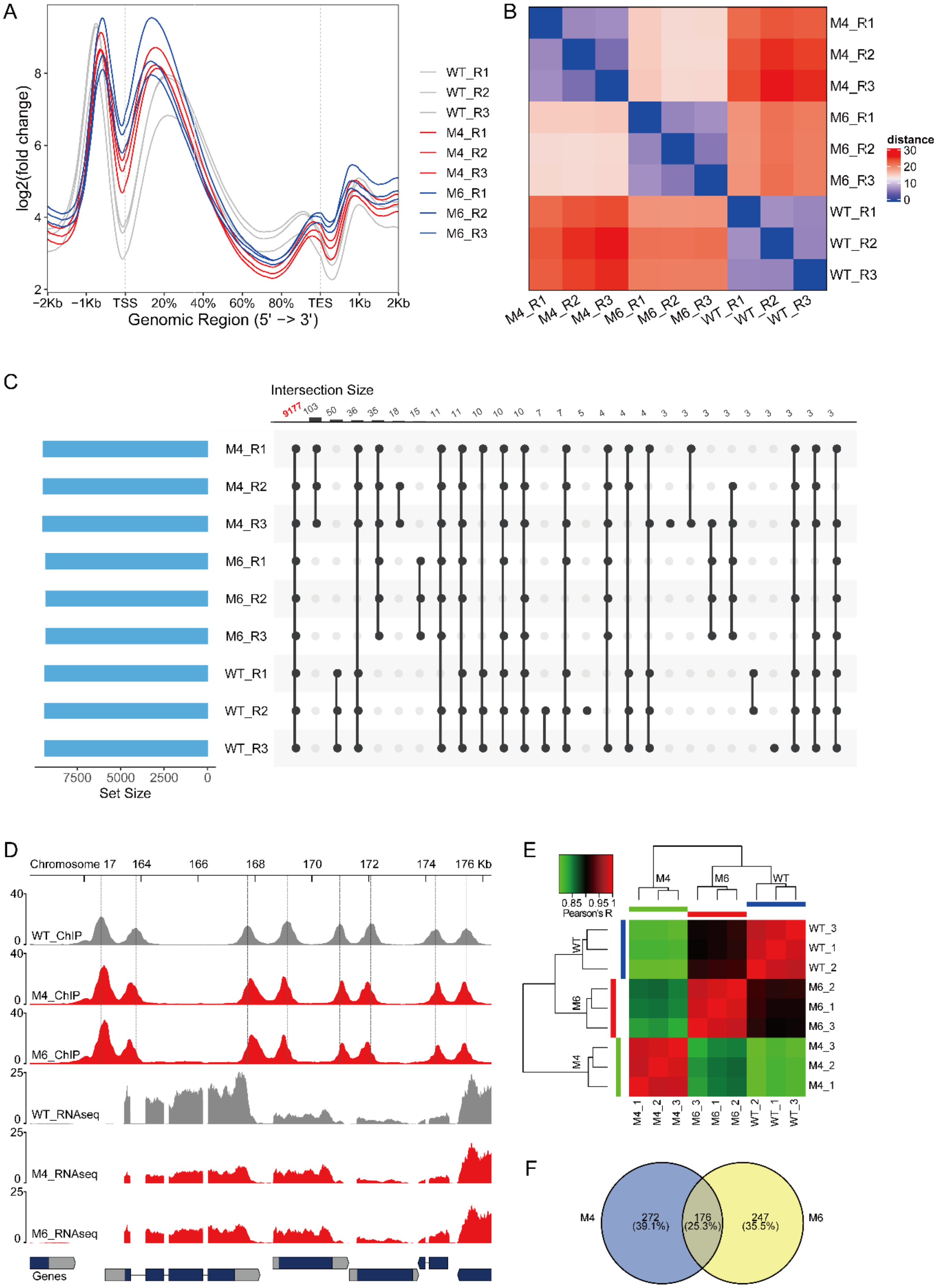


**Figure S7. Epigenetic and transcriptome profiling of NO24G02310-knockout mutants in *N. oceanica*.** (**A**) ChIP-Seq enrichment patterns for NO24G02310-knockout mutants. (**B**)

Distance matrix for different samples based on ChIP-Seq data. (**C**) Intersection of H3K4me2 peaks for different samples (NO24G02310-knockout mutants and wild-type). Items with < 2 shared peaks are not shown. (**D**) ChIP-Seq and RNA-Seq data tracks from different samples. The genomic region displays four pairs of H3K4me2 peaks. The vertical dashed lines indicate the positions of the ChIP-Seq peaks in the wild-type sample. (**E**) Heatmap and hierarchical clustering based on transcriptome similarities. The pairwise similarity between samples was computed as the Pearson correlation between the expression (TPM) of the differentially expressed genes. (**F**) Overlap of DEGs between M4 and M6 compared with WT.

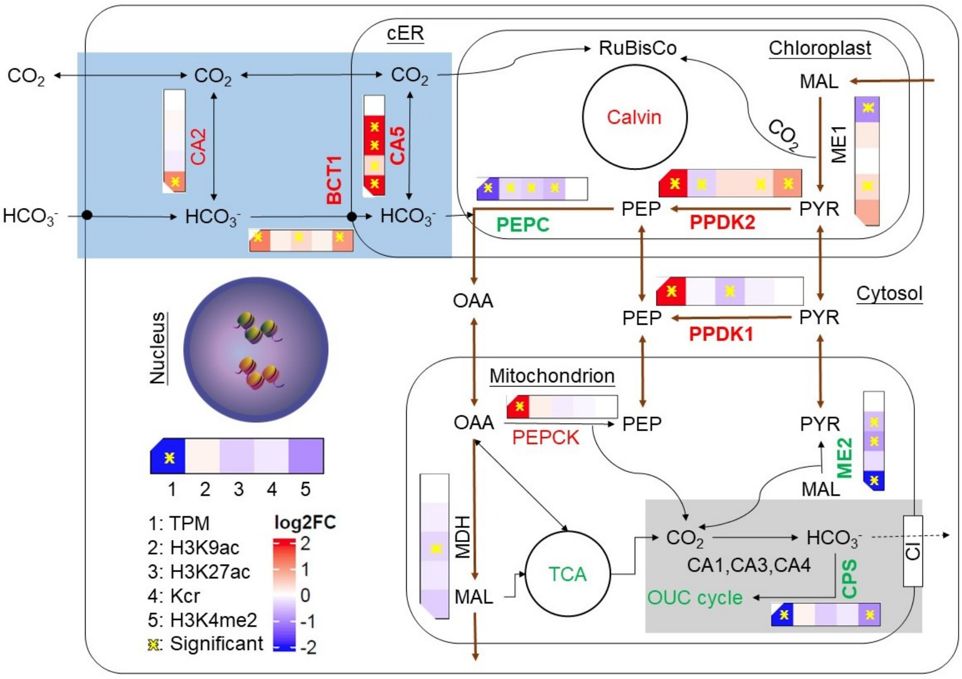


**Figure S8. The conceptual model of epigenetic regulation during CCM induction in *N. oceanica*.** CA, carbonic anhydrase; BCT, bicarbonate transporter; cER, chloroplast endoplasmic reticulum; PEPC, phosphoenolpyruvate carboxylase; PEPCK, phosphoenolpyruvate carboxylase kinase; ME, malic enzyme; OAA, oxaloacetate; CPS, carbamoyl phosphate synthase; CI: NADH-ubiquinone oxidoreductase complex I; PYR, pyruvate; MAL, malate; PPDK, pyruvate orthophosphate dikinase. Up-regulated genes (red), down-regulated genes (green), and differentially expressed genes with at least one type of epigenetic regulation (bold) are highlighted.
